# Efficient integration of spatial omics data for joint domain detection, matching, and alignment with stMSA

**DOI:** 10.1101/2024.07.29.605604

**Authors:** Han Shu, Jing Chen, Chang Xu, Jialu Hu, Yongtian Wang, Jiajie Peng, Qinghua Jiang, Xuequn Shang, Tao Wang

## Abstract

Spatial omics (SO) is a powerful methodology that enables the study of genes, proteins, and other molecular features within the spatial context of tissue architecture. With the growing availability of SO datasets, researchers are eager to extract biological insights from larger datasets for a more comprehensive understanding. However, existing approaches focus on batch effect correction, often neglecting complex biological patterns in tissue slices, complicating feature integration and posing challenges when combining transcriptomics with other omics layers. Here, we introduce stMSA (SpaTial Multi-Slice/omics Analysis), a deep graph contrastive learning model that incorporates graph auto-encoder techniques. stMSA is specifically designed to produce batch-corrected representations while retaining the distinct spatial patterns within each slice, considering both intra- and inter-batch relationships during integration. Extensive evaluations show that stMSA outperforms state-of-the-art methods in distinguishing tissue structures across diverse slices, even when faced with varying experimental protocols and sequencing technologies. Furthermore, stMSA effectively deciphers complex developmental trajectories by integrating spatial proteomics and transcriptomics data, and excels in cross-slice matching and alignment for 3D tissue reconstruction.

## 1 Introduction

Spatial omics (SO) technologies have greatly enhanced our understanding of molecular expression and interactions within tissue architectures [1]. These methodologies allow for the simultaneous analysis of thousands of genes, proteins, or metabolites, revealing their precise spatial distribution in tissues. Spatial transcriptomics (ST) is a leading technique in this field, providing detailed views of the transcriptome in spatial perspective [2]. Recently, spatial CITE-seq (Cellular Indexing of Transcriptomes and Epitopes by Sequencing) [3] has emerged as a significant advancement in spatial omics. This technique integrates RNA sequencing with the measurement of surface protein expression using specific antibodies, thereby enriching transcriptomic data with valuable spatially-resolved proteomic information. This integration allows for a comprehensive understanding of the spatial distribution of proteins, offering invaluable insights into the proteomic landscape within tissues. These advancements have paved the way for novel applications in understanding cellular interactions, identifying spatial biomarkers, and unraveling complex developmental and disease processes [4].

In tandem with the evolution of SO technologies, numerous computational methods have been proposed to analyze individual slices. A critical step in these analyses is the identification of spatial domains, which involves clustering spots with similar molecular expression profiles and spatial locations in an unsupervised manner. Accurate integration of molecular expression and spatial information is essential for this process. For instance, STAGATE [5] utilizes self-attention mechanisms and cell type-aware spatial graphs to learn latent representations of spots, while CCST [6] employs a Deep Graph Infomax (DGI) model, providing a general framework for multiple ST analysis tasks, including spatial domain detection and cell cycle identification. Although these methods have successfully analyzed individual slices, they remain limited to two-dimensional (2D) analyses due to their single-slice focus.

As the availability of SO data derived from the same tissues, organs, or embryos increases, there is a growing need to integrate multiple slices [7]. When slices are oriented horizontally, this integration facilitates extended spatial analyses; when vertical, it enables the reconstruction of three-dimensional (3D) models that more accurately reflect true spatial structures and cellular functions. While some methods, such as Scanorama [8] and Harmony [9], have been adapted for integrating SO data, their performance is often limited due to inadequate utilization of spatial information [10]. Methods like SEDR [11] address batch effects by learning latent representations for each slice using deep autoencoders; however, they overlook cross-slice relationships. Recently proposed methods, such as STAligner [12] and STitch3D [13], specifically aim to integrate multiple SO slices but still encounter limitations. For instance, STAligner employs mutual nearest neighbors (MNNs) for integration, while STitch3D requires additional reference data that can impact performance. Moreover, there are currently no effective methods for integrating spatial multi-omics data, such as combining transcriptomics with proteomics, to reveal intricate biological patterns from both perspectives.

To address these challenges, we present stMSA, a novel graph contrastive learning-based method specifically designed for integrating SO data from multiple sources. stMSA employs a contrastive learning strategy, optimizing two key principles to simultaneously learn the embeddings of cross-batch spots. First, within each slice, the embeddings of spots in connected adjacent microenvironments are made similar, while those of randomly sampled spots remain dissimilar. Second, embeddings of cross-batch spots with similar gene expression profiles are optimized to be similar, while those of randomly sampled cross-batch spots diverge. By adhering to these principles, stMSA effectively captures both inner-slice and cross-slice distribution patterns. We conducted a comprehensive evaluation of stMSA’s performance across various downstream tasks, including batch-effect correction, joint domain detection, cross-slice matching, and multi-slice alignment for 3D reconstruction. Our findings demonstrate that stMSA effectively eliminates batch effects and integrates data across diverse sequencing platforms. Additionally, stMSA successfully deciphers the developmental trajectory of B cells by integrating spatial proteomics and transcriptomics data. Overall, stMSA offers a robust and comprehensive solution for integrating multiple SO slices, addressing the limitations of existing methods, and facilitating advanced analysis of spatial transcriptomics and multi-omics data from various sources.

## 2 Materials and Methods

### 2.1 Data pre-processing

To prepare the data for analysis, the raw gene expressions undergo an initial log transformation and normalization using the Python package SCANPY [14]. To overcome the zero inflation in spatial transcriptomics data, similar to STAGATE, stMSA selects the top *k* highly variable genes (HVGs) as input features. For SO slices with common gene backgrounds, such as the DLPFC dataset [15], we set *k* as 3,000 by default. For SO slices with varying genes, usually derived from different platforms, it is suggested to set *k* higher and use the intersecting genes as input features. For proteomics and transcriptomics integration, we use the Principal Component Analysis (PCA) [16] to decompose the protein and RNA data and use the top 300 PCs as the input features.

### 2.2 Spatial graph construction for SO slices

We first construct a spatial graph for each slice based on the spot coordinates. Given the spot coordinates on slice *t*, represented as *s*_*t*_, we denote the spatial information of all *k* slices as set *S* = {*s*_1_, *s*_2_, …, *s*_*k*_}. stMSA initially calculates the Euclidean distances between all spots within each slice. Then stMSA uses a radius threshold *r* to define the edges between spots. In our experimental settings, we fine-tuned the radius threshold *r* to achieve spatial graphs with an approximate connectivity of 6 neighbors for each spot. The adjacency matrices for spatial graphs are denoted as 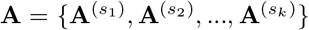 where 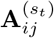 is set as 1 only if spot *i* and spot *j* within slice *s*_*t*_ are connected, and it is assigned 0 otherwise. For the unbalanced datasets, we randomly sampled the large slice into two small datasets and treated them as two individual slices.

### 2.3 Representation learning via parameter-shared graph auto-encoder

In order to effectively couple gene expression and spatial information, stMSA uses the technique of parameter-shared graph autoencoder to learn the latent embeddings of spots. Specifically, the graph auto-encoder consists of a graph attention network (GAT)[17] and a fully connected network (FCN)[18] for both the encoder and the decoder. Furthermore, to facilitate model convergence, stMSA shares the GAT and FCN parameters between the encoder and the decoder.

#### 2.3.1 Encoder

stMSA adopts the GAT network to aggregate spot features within a local microenvironment on a slice. In contrast to graph convolutional neural networks, GAT uses the self-attention mechanism to dynamically compute the importance of inter-spot connections during the feature aggregation process. Specifically, the GAT layer takes the gene expression profiles and the spatial graph of each slice as inputs. Suppose the gene expression profiles on slice *s*_*m*_ are denoted as 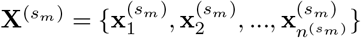 where 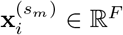 indicates the gene expression of spot *i*, and 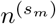 denotes the number of spots in slice *s*_*m*_, the formulation of the GAT layer is described as follows:

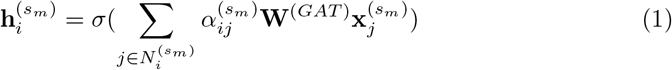

where 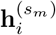 denotes the initial latent embedding of spot *i* within slice *s*_*m*_, *σ*(*·*) represents the aggregation function applied to neighboring spots. Specifically, the aggregation method selected in stMSA is the summation. 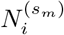 denotes the set of neighbors for spot *i* within slice *s*_*m*_. **W**^(*GAT*)^ denotes the trainable weight matrix, while 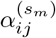 denotes the attention coefficient between spot *i* and its neighbor spot *j* within slice *s*_*m*_, computed via the following formula:

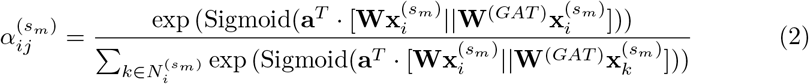

where **a**^*T*^ denotes the trainable attention coefficient vector responsible for weighting the closeness between the centroid spot and its neighboring spots. The Sigmoid(*·*) is employed as an activation function. Additionally, [*·*||*·*] denotes the concatenation operation.

Subsequently, stMSA employs a fully connected network further to compress the initial latent embedding into a lower dimension as shown in the following equation:

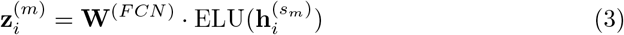

where 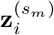 denotes the latent representation learned by stMSA, **W**^(*FCN*)^ denotes the trainable weight matrix, and ELU(*·*) denotes the Exponential Linear Unit activation function [19].

#### 2.3.2 Decoder

The decoder in stMSA also consists of a GAT layer and an FCN layer, both designed to reconstruct the input gene expressions. To expedite convergence and enhance efficiency, the decoder layers share the parameters learned from the encoder layers. Specifically, stMSA utilizes the transposed weight matrix of the encoder as the weight matrix for the corresponding decoder layers. Additionally, it directly incorporates the attention coefficients from the encoder’s GAT layer into the decoder’s GAT layer. The formula for the decoder is as follows:

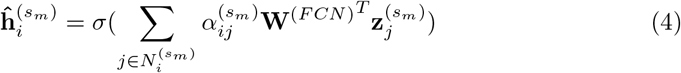

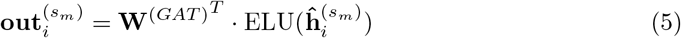

where 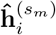 represents the reconstructed representation generated by the GAT decoder, while 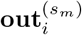 denotes the output of the reconstructed gene expression profiles.

### 2.4 Enhancing multi-slice integration via graph contrastive learning

To create a unified embedding space for spot features extracted from various slices, stMSA employs a contrastive learning approach to remove batch effects while preserving the original spatial patterns of the spots. In contrast to conventional graph contrastive learning methods that require building a corrupted graph, our approach leverages spatial graphs constructed from multiple input SO slices. It consists of two components: inner-batch contrastive learning and cross-batch contrastive learning.

#### 2.4.1 Inner-batch contrastive learning

In the inner-batch contrastive learning stage, the model is constrained to generate embeddings that closely resemble those of adjacent spots, and separate those non-adjacent spots. To achieve this, stMSA maximizes the embedding similarity between a centroid spot *i* and a spot in its adjacent micro-environment (denoted as positive pairs), while minimizing its similarity with a randomly sampled spot *k* (denoted as negative pairs) within the same batch (i.e. slice), and this process is formulated as follows:

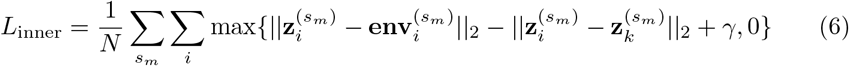

where *N* denotes the total number of spots across all slices, 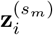 represents the embedding of the centroid spot *i* on slice *s*_*m*_, 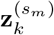 refers to the embedding of the randomly sampled spot, *γ* is the margin parameter, typically set to 1.0 by default. The 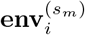 denotes the micro-environment embedding constructed from the embeddings of neighboring spots of spot *i*, which is calculated as follows:

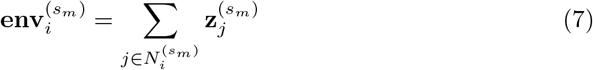

#### 2.4.2 Cross-batch contrastive learning

In the cross-batch contrastive learning stage, stMSA aims to generate batch-corrected embedding for each spot, which also reserves the spatial gene expression pattern. To achieve this, stMSA maximizes the embedding similarity between the centroid spot *i* and a spot *j* with similar gene expression in another slice (denoted as positive pairs), while simultaneously minimizing the similarity with a randomly sampled spot *k* (denoted as negative pairs) across slices. This process is depicted as follows:

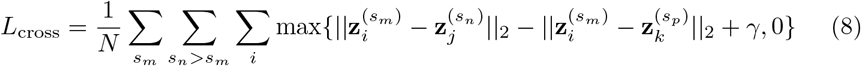

where 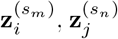 and 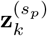 represent the embeddings of centroid spot *i* in slice *s*_*m*_, its positive pair mate *j* in slice *s*_*n*_, and its negative pair mate *k* in slice *s*_*p*_, respectively. The positive pairs are selected only when spot *j* is the most similar spot to spot *i*, and spot *i* is the most similar spot to spot *j* simultaneously.

The objective function of the two contrastive learning stages can be expressed as follows:

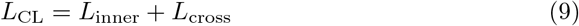

Due to the large number of spots in each batch and the relatively low neighbor count that we have set, the probability of sampling nearby spots in randomly selected negative pairs is extremely low. This ensures that the model can effectively discriminate the similarities and differences among spots. Additionally, incorporating the minimization of similarity between negative pairs helps prevent a scenario in which the embeddings of all spots become excessively similar, which will affect the performance of downstream tasks.

### 2.5 Gene expression reconstruction via graph autoencoder

The goal of gene expression reconstruction (GER) is to generate embeddings that can capture unique gene expression patterns, preserving their inherent characteristics while avoiding excessive similarity to neighboring spots. To accomplish this, the objective is to maximize the similarity between the input gene expression and the reconstructed output. The loss function is expressed as follows:

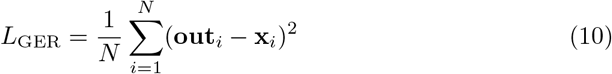

Where *N* represents the total number of spots in the input data, **out**_*i*_ and **x**_*i*_ represent the reconstructed gene expression and input gene expression of spot *i*, respectively.

### 2.6 Domain distribution tuning with deep embedding clustering

The domain distribution tuning (DDT) task aims to refine the clustering performance using the deep embedding clustering (DEC) technique[20]. The primary goal is to minimize the distance between the soft clustering distribution[21] and the auxiliary target distribution, as depicted below:

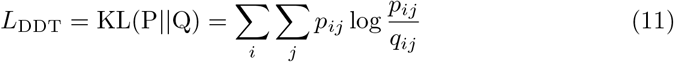

Where *Q* and *P* denote the soft clustering distribution and the auxiliary target distribution, respectively. *p*_*ij*_ and *q*_*ij*_ represent the probability that a spot *i* belongs to a cluster *j* in distribution *P* and *Q*, respectively. The soft clustering distribution and the auxiliary target distribution are computed as follows:

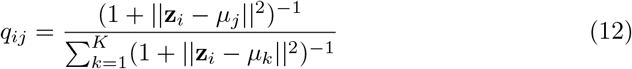

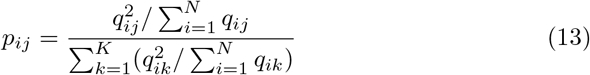

Where *µ*_*j*_ denotes the initial cluster centroid of a cluster *j*, and *K* represents the cluster count. The initial cluster centroids and cluster count are generated using Louvain clustering[22], which takes gene expression as input and we set the Louvain resolution parameter to 1.0 by default.

In sum, the overall loss function is defined as follows:

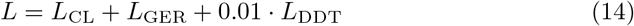

### 2.7 Joint domain detection across SO slices

After encoding the spot features across different SO slices into a coherent embedding space, we use the unsupervised clustering technique to identify the spatial domains in the joint embedding space. In particular, for datasets with known labels or prior knowledge of the cluster number, stMSA employs the finite Gaussian mixture clustering model from the R package MCLUST [23] to identify spatial domains. In cases where the cluster number is unknown, stMSA utilizes the Louvain clustering algorithm for community detection [22] which has been implemented in the Python package SCANPY [14].

To evaluate the performance of domain detection, we utilize the metric of adjusted Rand index (ARI) [24], which is calculated using the following formula:

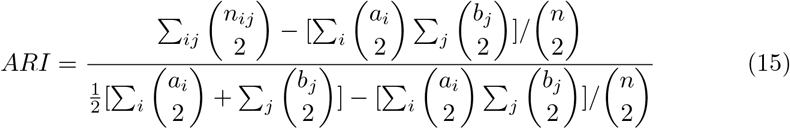

where *n*_*ij*_ denotes the count of spots within the predicted cluster *i* that are assigned to the ground truth class *j, a*_*i*_ represents the count of spots within the predicted cluster *i, b*_*j*_ denotes the count of spots within the ground truth class *j*, and *n* represents the total count of spots. A higher ARI score indicates a more accurate detection of the domains that align with the ground truth labels. The ARI score was calculated using the Python package Scikit-learn [25].

To evaluate the spatial autocorrelation of the learned latent representations, we utilize the Geary’s *C* [26], which is calculated as follows:

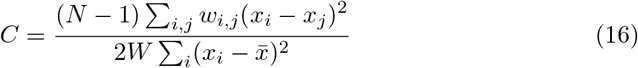

where *N* denotes the number of spots, *x*_*i*_ and *x*_*j*_ denote the latent representation of spot *i* and *j, w*_*ij*_ denotes the similarity of spot *i* and *j*, and *W* is the sum of all the similarity. A lower Geary’s *C* score indicates a higher degree of spatial autocorrelation.

The Moran’s *I* [27] is also a spatial autocorrelation metric we use to evaluate the latent representation. The Moran’s *I* can be calculated as follows:

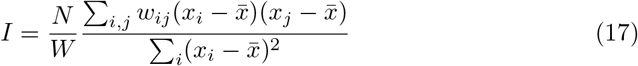

where *N, x*_*i*_, *x*_*j*_, *w*_*ij*_, and *W* retain their previous meanings. A higher Moran’s *I* score suggests a greater level of spatial autocorrelation. The Geary’s *C* and Moran’s *I* score are calculated by the Python package SCANPY [14].

### 2.8 Cross-slice matching

The objective of matching spots across two adjacent SO slices is to establish a mapping between pairs of spots from different slices, ensuring that the matched spots exhibit similar gene expression patterns. Additionally, the mapping should preserve the spatial relationship of spots within each SO slice. To achieve this, we calculate the Euclidean distances in the embedding space between each spot *s* on the source slice and all other spots on the target slice. The spot *t* on the target slice with the smallest Euclidean distance to spot *s* is considered its match. To evaluate the quality of the matching, we compare the identity of the spot’s label (such as cell type, tissue type) for each pair, and calculate a matching score *acc* as depicted below:

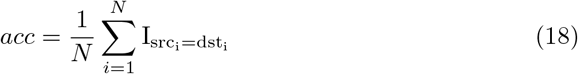

Where *src*_*i*_ and *dst*_*i*_ represent the cell types of spot *i* in the source slice and its corresponding spot in the target slice, respectively. The indicator function 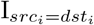 evaluates to 1 if *src*_*i*_ and *dst*_*i*_ represent the same cell type, and 0 otherwise. The *acc* score quantifies the accuracy of the matching process. We utilize the Python package FAISS[28] to improve the efficiency of matching spots.

### 2.9 Multi-slice alignment

Aligning multi-slice data is a key step in reconstructing 2D tissue slices into 3D. In this regard, stMSA incorporates landmark points to guide the alignment process. Given the absence of dedicated SO alignment methods, we have enhanced the Iterative Closest Point (ICP) algorithm [29] to align slices using the domain detection output from stMSA. Furthermore, we have introduced a scoring method named Ali-score to evaluate the quality of the alignment result.

#### 2.9.1 ICP with disturbance

The alignment procedure employed in stMSA comprises three primary components: (1) integration of multi-slice data using stMSA, (2) identification of the landmark domain, and (3) derivation of the transformation matrix through a modified version of the Iterative Closest Point (ICP) algorithm.

The ICP algorithm is renowned for its efficiency and simplicity, making it one of the most extensively utilized point cloud registration methods. The objective function for the ICP alignment is stated as follows:

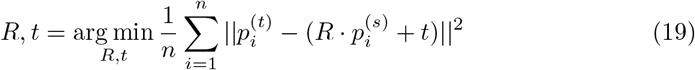

where *R* and *t* denote the rotation matrix and translation vector, respectively. 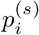 and 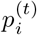 represent the source and target point clouds, respectively, and *n* denotes the point count.

Given the non-convex nature of the problem and the dependency on local iterative steps, the ICP algorithm is prone to converging to local minima [30]. To address this concern, stMSA incorporates a threshold, where the mean error between two iterations should be below the threshold to halt the iteration process. If the threshold is surpassed, the point cloud undergoes rotation, and the iteration persists. Empirically, this strategy effectively enables ICP to avoid local minima.

#### 2.9.2 Alignment scoring method

Currently, there is a lack of a standardized metric for evaluating the alignment results of multi-slice data. In this study, we introduce a novel scoring system called Ali-score, specifically designed for assessing the quality of alignment. The goal of multi-slice alignment is to align the positions of different slices to a global coordinate system, ensuring that cells within the same domain are accurately positioned in corresponding 3D regions across slices.

To evaluate the effectiveness of this alignment method, we begin by identifying the nearest spot in the slice below through the calculation of the Euclidean distance between their aligned coordinates. In order to assess the alignment outcome, we compare the predicted cell type of each spot pair and calculate the percentage of correctly matched spot pairs using the following approach:

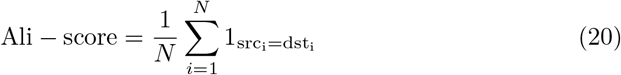

The Ali-score serves as a quantitative measure of alignment accuracy, where a higher Ali-score signifies a closer proximity between spots of the same cell type within the aligned regions, indicating a more effective alignment. To calculate the Ali-score, we have developed a customized Python function. Moreover, our algorithm supports the identification of the *k* nearest neighbors for each spot and incorporates a voting mechanism based on domain labels, allowing for the computation of a more flexible Ali-score.

### 2.10 The overall structure of stMSA

The dimensions of the layers in the auto-encoder are set as follows: the input data shape is 512 in the encoder GAT layer, and 512 to 30 in the encoder FCN layer. The decoder is symmetric to the encoder. To optimize the model parameters, we use the Adam optimizer [31] with a learning rate of 0.001. We implement the stMSA model using the popular graph neural network framework PyTorch Geometric (PyG) [32]. We set the number of training epochs to 500. In the cross-batch SMR task, we update the cross-batch pairs, and for the DDT task, we update the DEC p distribution every 100 epochs.

### 2.11 Benchmarking

The performance of stMSA in multi-slice representation learning, integration, and domain identification is compared with seven state-of-the-art methods: Scanorama [8], Harmony [9], STAGATE [5], SEDR [11], STAligner [12], Stitch3D [13], and SPACEL [33]. Additionally, the cross-slice matching capabilities of stMSA are assessed against SLAT [34], while its alignment performance is compared specifically with STAligner [12]. Details on the methods compared are available in Supplementary Note S1.

To evaluate the performance of integration, we calculate the Local Inverse Simpson’s Index (LISI)[9] as follows:

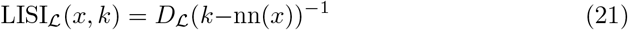

where *x* represents the set of object embedding vectors, *L* denotes the corresponding label set for the vector set, *k* is a perplexity parameter set to 90 by default, *k*−nn(*·*) represents the k-nearest neighbors algorithm, and *D*_*L*_(*·*) denotes Simpson’s index, computed as follows:

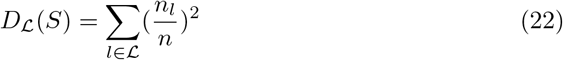

where *S* denotes the object vector set, *n*_*l*_ represents the count of vectors belonging to label type *l*, and *n* is the overall count of vectors in set *S*.

The LISI score serves as an indicator of the degree of mixing between different label types, where a higher LISI score signifies improved cohesion and enhanced elimination of batch effects. The LISI score was calculated using the Python package HarmonyPy[9].

To evaluate domain detection, we use the Adjusted Rand Index (ARI) [24], which measures the congruence between the ground truth labels and the predicted clustering results. Further details on ARI are provided in the Multi-slice domain detection section. Moreover, we employ Geary’s *C* [26] and Moran’s *I* [27] to assess the spatial autocorrelation of the latent representation. The matching results of cross-slice matching are assessed using a Matching Score, detailed in the Cross-slice matching section. Finally, the alignment results are evaluated using the Ali-score, with specifics available in the Alignment scoring method section.

### 2.12 Data availability

Multiple SO datasets derived from human and mouse tissues, organs, and embryos are used to evaluate the performance of stMSA. Specifically, the DLPFC dataset can be obtained from the Lieber Institute GitHub repository at https://github.com/LieberInstitute/HumanPilot. The mouse brain serial dataset can be accessed at STOmicsDB [35] (Anterior: https://db.cngb.org/stomics/datasets/STDS0000018, Posterior: https://db.cngb.org/stomics/datasets/STDS0000021). The mouse brain coronal Frozen and DAPI dataset can be found at squidpy dataset (Frozen: https://squidpy.readthedocs.io/en/stable/api/squidpy.datasets.visium_hne_adata.html, DAPI: https://squidpy.readthedocs.io/en/stable/api/squidpy.datasets.visium_fluo_adata.html). The mouse brain coronal FFPE dataset can be found at STOmicsDB (https://db.cngb.org/stomics/datasets/STDS0000052). The Slide-seq V2 mouse brain bulb data can be obtained from the Broad Institute webpage (https://singlecell.broadinstitute.org/singlecell/study/SCP815/highly-sensitive-spatial-transcriptomics-at-near-cellular-resolution-with-slide-seqv2#study-download). The Stereo-seq mouse olfactory bulb dataset can be accessed from the SEDR GitHub repository (https://github.com/JinmiaoChenLab/SEDR_analyses/tree/master/data).

The Visium mouse olfactory bulb can be found at GEO GSM4656181 dataset (https://www.ncbi.nlm.nih.gov/geo/query/acc.cgi?acc=GSM4656181). The Spatial Transcriptomics mouse olfactory bulb can be found at NCBI database BioProject ID PRJNA316587. The mouse organogenesis spatiotemporal transcriptomic atlas dataset can be found at STOmicsDB (https://db.cngb.org/stomics/mosta/spatial/). The Slide-seq V2 mouse embryo dataset can be found at cellxgene database (https://datasets.cellxgene.cziscience.com/acc80ff4-5dee-46cc-bf22-84a9a83c9c38.h5ad). The proteomics human tonsil dataset can be found at GEO GSE213264 dataset (https://www.ncbi.nlm.nih.gov/geo/query/acc.cgi?acc=GSE213264). The 10X Visium human tonsil dataset can be found at 10X Genomics databse (https://www.10xgenomics.com/datasets/visium-cytassist-gene-and-protein-expression-library-of-human-tonsil-with-add-on-antibodies-h-e-6-5-mm-ffpe-2-standard). The mouse brain coronal dataset can be found at STOmicsDB (https://db.cngb.org/stomics/datasets/STDS0000218). A summary of the dataset information can be found in Supplementary Table S1.

## 3 Results

### 3.1 Overview of stMSA

This study introduces stMSA, a novel deep graph representation learning model designed for integrating multiple spatial omics data (Fig. 1A). The primary objective of stMSA is to generate consistent cross-batch spot embeddings, essential for tasks such as joint domain detection, cross-slice matching, and 3D reconstruction alignment (Fig. 1B).

**Fig. 1.**
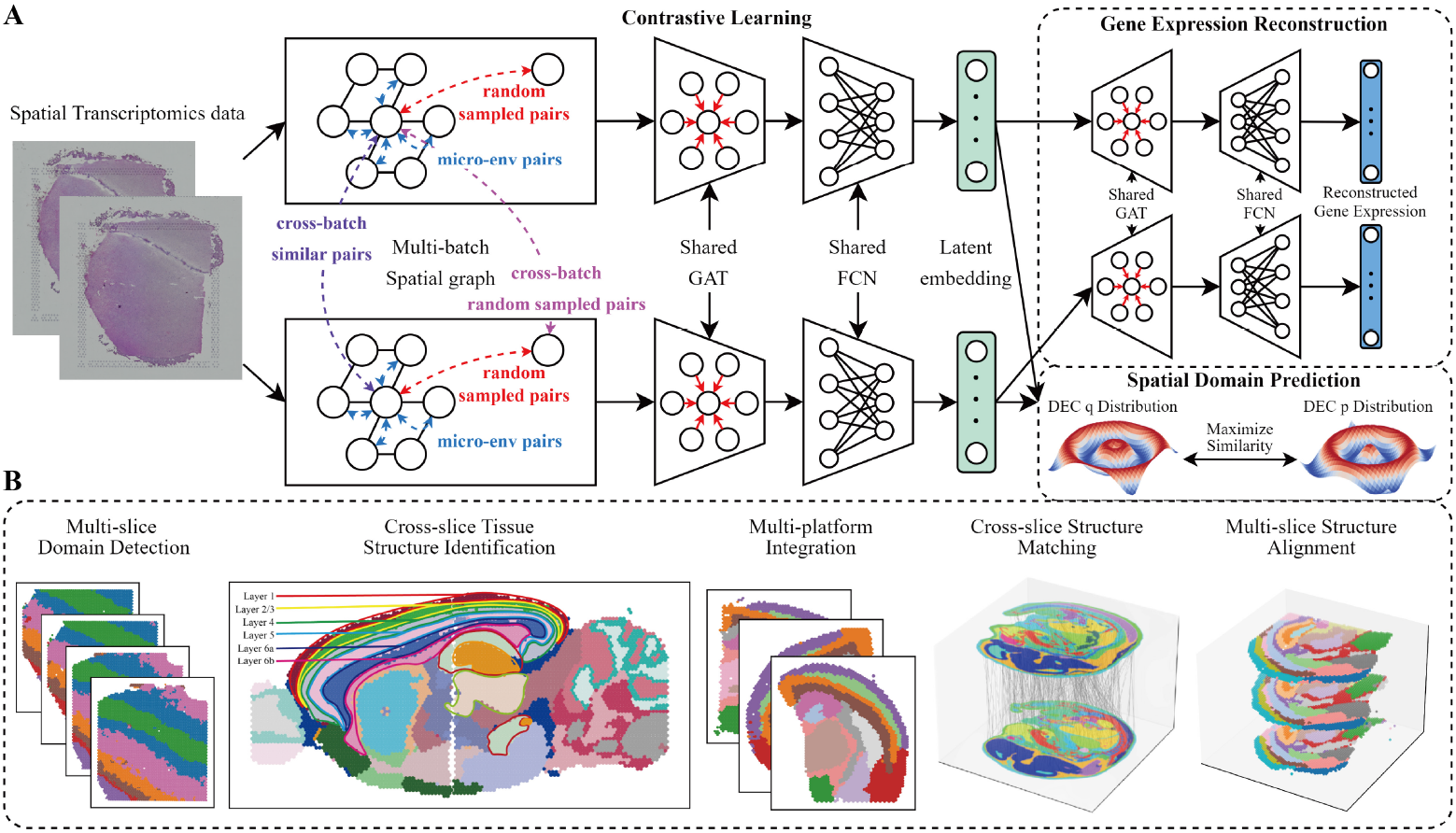
Overview of the stMSA framework. **(A)** stMSA leverages multi-slice omics expression and spatial coordination information as input. It utilizes an auto-encoder to learn latent batch-corrected multi-slice representation and optimizes the model through three distinct optimization tasks. **(B)** The latent embedding learned by stMSA serves various downstream tasks, including multi-slice domain detection, cross-slice tissue structure identification, multi-platform integration, cross-slice matching, and multi-slice alignment.

The model processes pre-processed omics expression data and spatial information from multiple slices as input. For each slice, stMSA constructs a spatial graph using the coordinates of each spot. It then utilizes both the omics expression profiles and the spatial graphs to derive latent representations. The core of stMSA is a contrastive learning strategy that simultaneously considers the inner-batch patterns and the cross-batch patterns to integrate data across multiple slices. This method aims to maximize the similarity of spatially adjacent spots within the same slice and across different slices that exhibit similar gene expression patterns, while minimizing the similarity between dissimilar spot pairs, both within and between slices.

In detail, stMSA enhances the similarity between centroid spots and their associated micro-environmental latent representations, enabling the model to capture detailed local features from neighboring spots. To preserve and highlight the unique expression patterns of each spot, the model also optimizes gene expression reconstruction through an attention-based graph auto-encoder, a process we term the gene expression reconstruction (GER) task. Furthermore, we refine the model’s performance by dynamically optimizing the predicted spatial domains using the deep embedding clustering (DEC) method [20].

### 3.2 stMSA enhances joint domain identification for human dorsolateral prefrontal cortex slices

We applied stMSA to the human dorsolateral prefrontal cortex (DLPFC) dataset from the 10x Visium platform [15], which includes 12 slices from three donors with detailed spatial annotations (Fig. 2A). To assess its performance in domain identification, we compared stMSA against seven state-of-the-art methods: Scanorama [8], Harmony [9], STAGATE [5], SEDR [11], STAligner [12], Stitch3D [13], and SPACEL [33]. Domain detection performance was evaluated using the Adjusted Rand Index (ARI) [24] for each method.

**Fig. 2.**
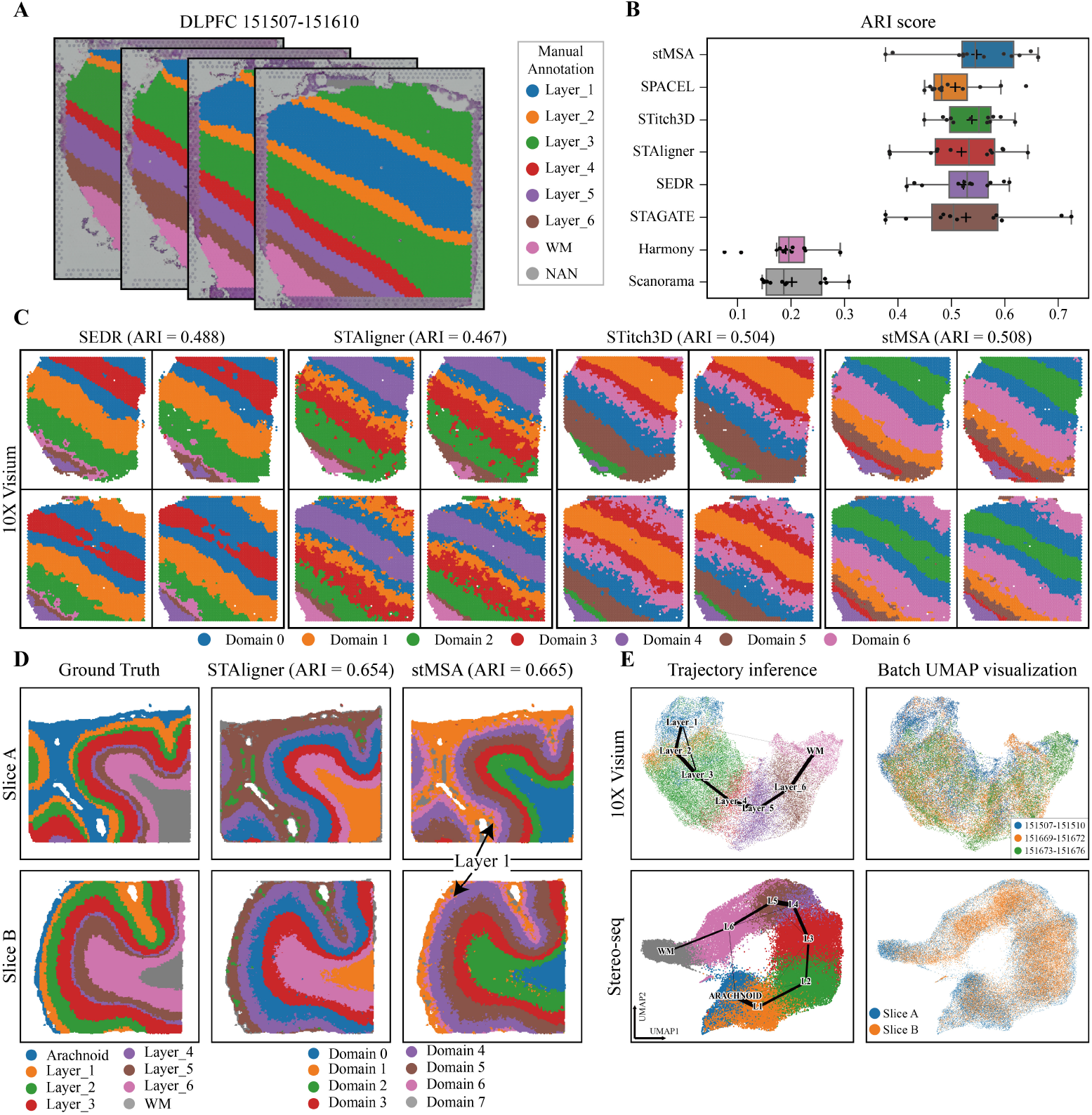
stMSA improves joint domain identification for human dorsolateral prefrontal cortex slices. **(A)** Manual annotations, ranging from the cortex layer 1 to the white matter, are provided with the histology image of a donor (slice identifiers: 151507-151610). **(B)** The box plot illustrates the ARI scores across 12 10X Visium DLPFC slices, generated by training separately for each donor. **(C)** Visualization of domain detection results for SEDR, STAligner, SPACEL, and stMSA. The domain detection processes are conducted using all four slices as inputs. **(D)** The ground truth, STAligner, and stMSA domain detection result of the two stereo-seq obtained DLPFC slices. **(E)** The trajectory inference plot generated based on the embeddings learned by stMSA using all 12 DLPFC slices for the 10X Visium and the two Stereo-seq DLPFC slices (The color reflects the ground truth domains). And the UMAP visualization for different donors/slices (The color reflects three donors/slices).

As shown in Fig. 2B, stMSA achieved an average ARI score of 0.544 across the 12 DLPFC slices, outperforming STitch3D (0.535) and SPACEL (0.508), both of which were significantly influenced by the quality of single-cell data. Visual comparison of spatial domain distributions for one donor (slices 151507-151610) further demonstrated stMSA’ superior performance, achieving an ARI score of 0.508 (Fig. 2C, Supplementary Fig. S2A). Other methods, like Scanorama (0.255) and Harmony (0.222), struggled due to a lack of spatial integration, while STAGATE (0.407) lacked multislice integration strategies. Despite moderate success from SEDR (0.488), Stitch3D (0.504), and SPACEL (0.436), these methods showed inaccuracies in layers 2-6, whereas stMSA produced more coherent boundaries and shapes. STAligner (0.467) was the closest competitor in spatial domain distribution but was still outperformed by stMSA in accuracy and boundary clarity.

To assess the scalability of stMSA, we applied it to two DLPFC slices from the Stereo-seq platform [36], each containing over 30,000 spots (Fig. 2D, Supplementary Fig. S2B). Here, stMSA and STAligner accurately represented the layer structures, while SEDR failed to capture the zonal domains. Trajectory analyses [37] confirmed that stMSA effectively traced the path from white matter to the arachnoid membrane (Fig. 2E), unlike SEDR, which conflated multiple layers. Notably, stMSA distinguished the thin Layer 1 (domain 6) from the adjacent layers, achieving the highest ARI score of 0.665 across both slices.

We assessed stMSA’ effectiveness in removing batch effects using UMAP visualizations and LISI (Local Inverse Simpson’s Index) batch mixture score (Supplementary Fig. S2B, C, and S3). The embeddings from three donors were well mixed, with no clear batch effects observed in the 10X Visium dataset. This trend continued in the Stereo-seq dataset, confirming successful integration across donors. Additionally, we evaluated computational efficiency. stMSA and STAGATE processed up to 20 slices with less than 12 GB of GPU memory, while Stitch3D and SEDR struggled to handle larger datasets (Supplementary Fig. S4).

### 3.3 Identification of cross-batch structures using stMSA in multi-slice mouse brain

We applied stMSA to a sagittal section dataset of the mouse brain obtained via 10x Visium sequencing, which includes multiple slices due to the large tissue volume (Fig.3A). Each slice, covering up to 4,992 spots, was divided into anterior (Fig.3B, left) and posterior (Fig. 3B, right) sections for analysis. Our objective was to identify common spatial domains across slices and assign consistent domain labels. We compared the performance of stMSA with two effective methods, SEDR and STAligner, in this multi-slice dataset.

**Fig. 3.**
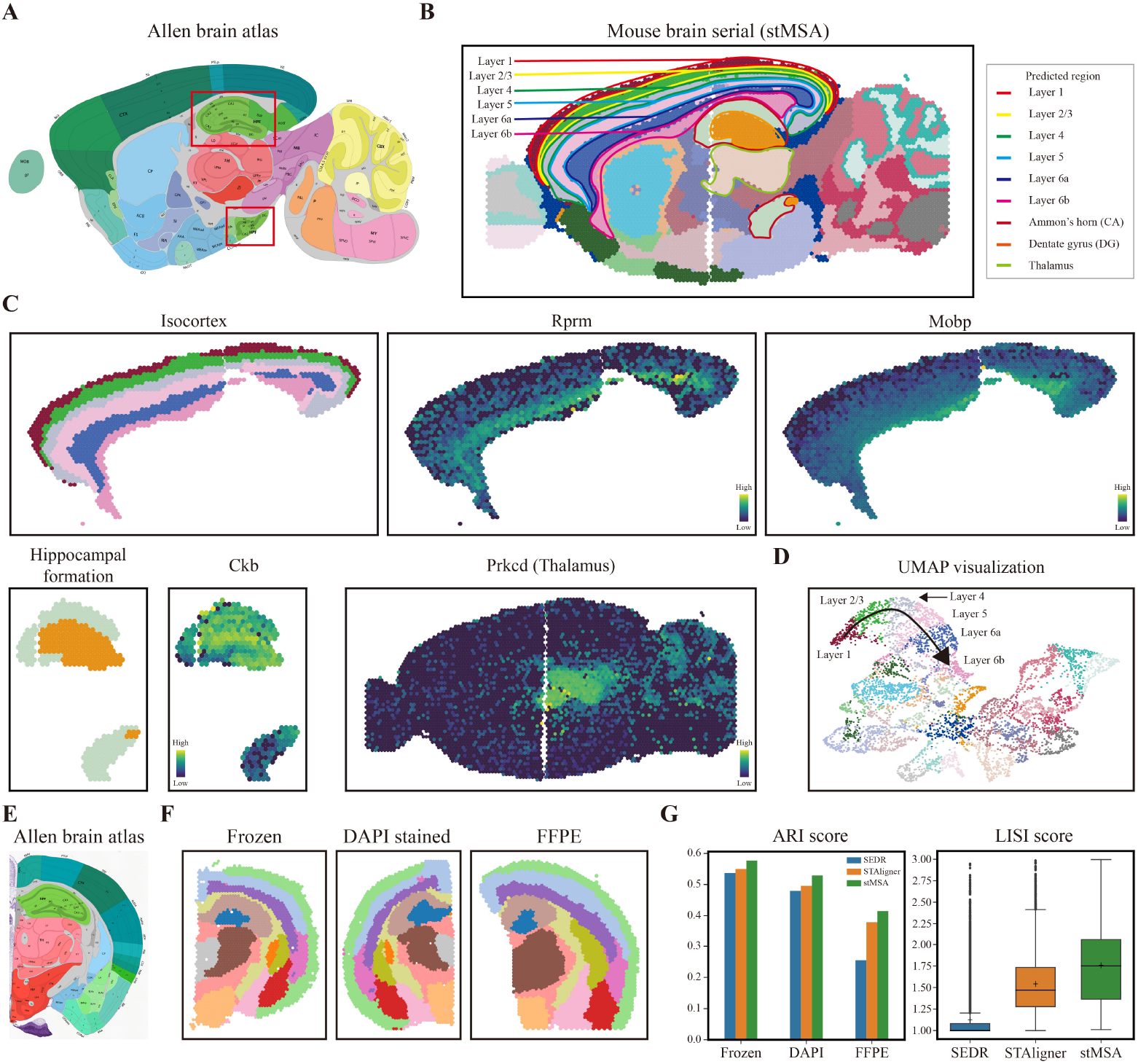
stMSA can identify cross-batch tissue structures in the 10x Visium-obtained mouse brain dataset. **(A)** The anatomical structure of sagittal mouse brain provided by the Allen Brain Atlas. **(B)** Spatial domains predicted by stMSA with manually annotated landmark domains (isocortex from layer 1 to layer 6b, hippocampus, and thalamus). **(C)** Subdomains of the isocortex, hippocampus, and thalamus, along with the expression levels of their corresponding marker genes. **(D)** The UMAP visualization of latent embeddings derived by stMSA. The substructures of the isocortex from layer 1 to layer 6b are manually annotated on the plot. **(E)** The anatomical structure of coronal mouse brain provided by the Allen Brain Atlas. **(F)** The clustering result of stMSA in fresh frozen with H&E stained, DAPI stained and FFPE preserved coronal mouse brain slices. **(G)** The ARI clustering score and the LISI integration score for SEDR, STAligner, and stMSA in the three coronal mouse brain slices.

Fig.3B illustrates the domain detection performance of stMSA relative to the manually annotated structures from the Allen Brain Atlas (Fig.3A). Both stMSA and STAligner accurately represented cross-slice structures, including the six cortical layers in the isocortex and thalamus, while SEDR identified only five layers within the isocortex. Notably, stMSA uniquely identified the hippocampus and its inner subdomains, accurately delineating the “hook”-shaped outline of Ammon’s horn, whereas STAligner failed to distinguish this feature. SEDR inaccurately assigned the lower anterior section of the hippocampus to a domain that did not correspond to the expected structure across slices.

Upon further analysis of fine-grained structures, stMSA demonstrated the ability to accurately recognize subdomains within significant brain structures, such as those in the isocortex and hippocampus. Comparison of gene expression distributions with known markers confirmed the accuracy of these subdomain identifications (Fig. 3C). For example, genes like *Rprm* and *Mobp* exhibited consistent expression patterns corresponding to cortical layers [38–41], while *Ckb* showed a “curved tube” structure aligning with our hippocampal domain results [42]. The spatial distribution of the marker gene *Prkcd* for the thalamus also significantly overlapped with structures identified by stMSA. Additionally, UMAP visualizations and trajectory inference from embeddings learned by stMSA confirmed the correspondence between the relative positions of isocortex layers and their actual physical locations (Fig. 3D, Supplementary Fig. S5B).

Next, we assessed stMSA’s capability to integrate coronal mouse brain data from various experimental protocols, including fresh frozen tissue with H&E staining, fresh frozen tissue with DAPI staining, and formalin-fixed paraffin-embedded (FFPE) tissue with H&E staining (Supplementary Fig.S6A). We compared stMSA ‘ performance with SEDR and STAligner in analyzing these slices and obtaining embeddings in the same feature space, with domain predictions visualized separately in the slices (Fig. 4F, Supplementary Fig. S6B-D). Results indicated that both stMSA and STAligner consistently identified the same tissue structures across protocols, whereas SEDR struggled with consistency. For example, stMSA and STAligner assigned the same domain labels to the Dentate gyrus across all slices, with stMSA labeling it as domain 1 and STAligner as domain 8. In contrast, SEDR showed inconsistencies in tissue identification across protocols. Additionally, stMSA effectively distinguished between the hypothalamus and white matter, accurately labeling these structures in domain 7 and domain 15, respectively, while STAligner misclassified them into incorrect domains. stMSA achieved the highest domain detection ARI score across all slices, demonstrating its superior ability to learn high-quality latent representations. Furthermore, it recorded the highest average LISI scores, indicating enhanced capability to integrate data across slices and mitigate batch effects in the resulting latent representations (Fig. 4G).

**Fig. 4.**
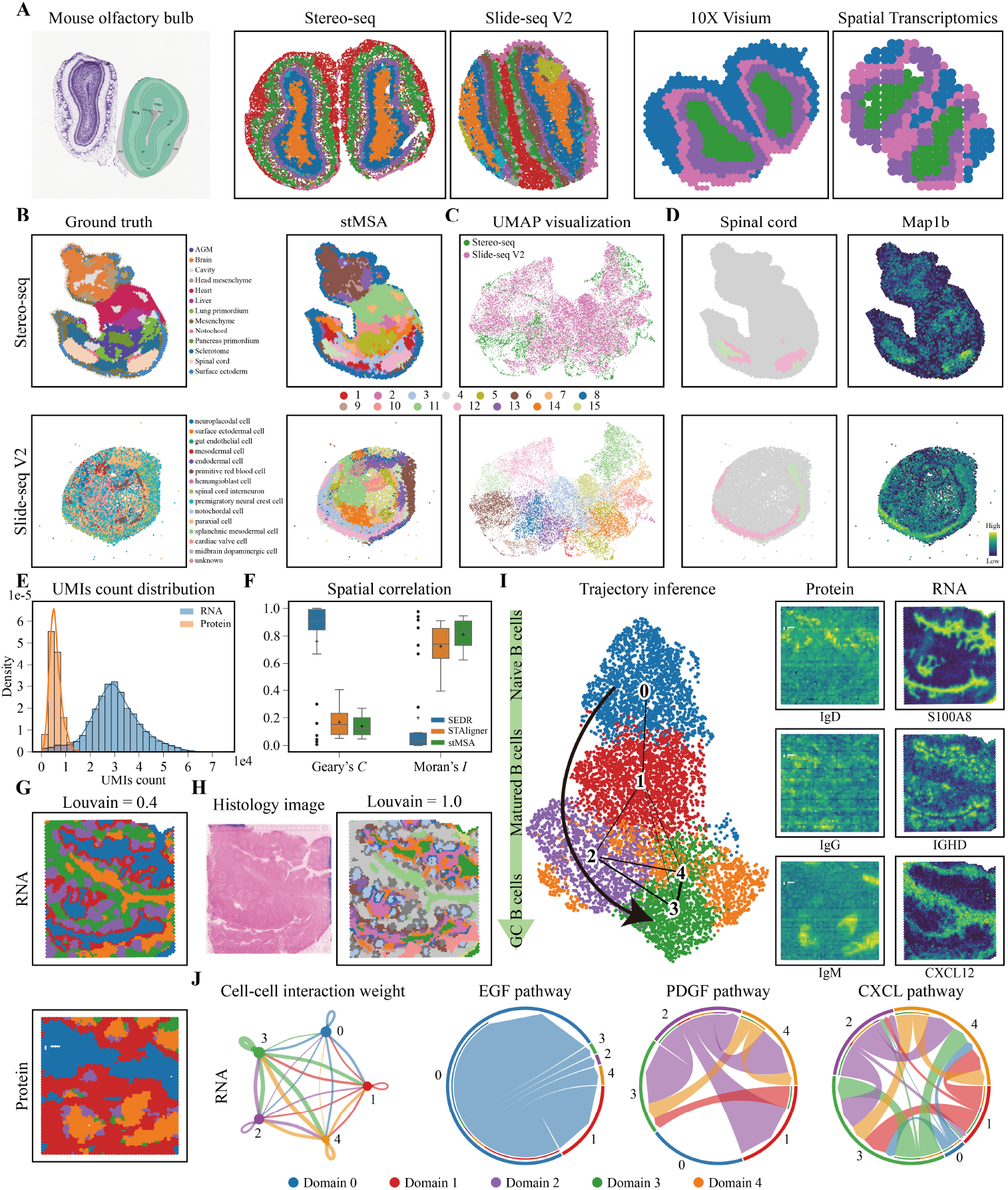
stMSA achieves robust performance across diverse sequencing platforms, different dataset size, and multi-omics data. **(A)** The allen brain atlas, the domain identification result for Stereo-seq and Slide-seq V2 platform and 10X Visium and Spatial Transcriptomics platform for mouse olfactory bulb slices. **(B)** The manual annotations and the domain identification result of stMSA in the Stereo-seq and Slide-seq V2 obtained mouse embryo dataset. **(C)** The UMAP visualization of the distribution of different sequencing platform and the clustering result. **(D)** The spatial distribution of spinal cord region stMSA predicted and its corresponding marker gene. **(E)** The unique molecular identifiers (UMI) count distribution for the protein and the RNA data in the human tonsil dataset. **(F)** The spatial correlation score Geary’s *C* and Moran’s *I* for the latent representation generated by SEDR, STAligner and stMSA. For Geary’s *C*, low *C* score indicated high correlation, and for Moran’s *I*, high *I* score indicated high correlation. **(G)** The domain identification result for the human tonsil protein and RNA data set louvain resolution to 0.4. **(H)** The histology image and the domain identification result for the human tonsil RNA data and set the resolution for louvain to 1.0. **(I)** The trajectory inference for the human tonsil dataset and the corresponding spatial distribution of the protein marker and gene marker for the naive B cells (IgD/S100A8), Matured B cells (IgG/IGHD), and GC B cells (IgM/CXCL12). **(J)** The cell-cell communication result and the gene pathway for the five domain stMSA identified.

Overall, our findings highlight stMSA’ effectiveness in identifying cross-batch tissue structures, accurately delineating subdomains, and integrating multi-slice datasets. This establishes stMSA as a powerful tool for analyzing complex brain architectures in spatial transcriptomics.

### 3.4 Robust integration of spatial omics data across sequencing platforms and dataset sizes

To evaluate the efficacy of stMSA in integrating spatial omics data from various sequencing platforms, we applied it to mouse olfactory bulb data acquired through both the Stereo-seq [43] and Slide-seq V2 [44] methods. Similar to the previous experimental settings, we jointly analyzed the two SO slices using each method, with domain prediction taking place in the same feature space. As depicted in Fig. 4A and Supplementary Fig. S7A, B, stMSA successfully recognized consistent structures in both slices, aligning well with the Allen Brain Atlas annotations. Supplementary Fig. S8A demonstrates that STAligner achieved similar results, while SEDR encountered difficulties in generating clear domain structures.

Next, we assessed stMSA’s performance with smaller datasets. Specifically, we applied stMSA to two mouse olfactory bulb slices obtained via the 10X Visium and Spatial Transcriptomics [45] platforms, which contained 918 and 267 spots, respectively. The domain identification results (Fig. 4A, Supplementary Fig. S7C) demonstrated that stMSA was the only method capable of accurately identifying the joint domain structure. In contrast, STAligner and SEDR incorrectly assigned the same domain in different slices while using distinct cluster labels. Additionally, UMAP visualizations (Supplementary Fig. S7D) indicated that stMSA effectively mitigated batch effects across the two sequencing platforms, validating its capability to learn a batch-effect-free latent representation even in small-scale datasets.

Furthermore, we examined stMSA’s ability to integrate data from unbalanced dataset sizes, as different sequencing platforms may capture data at varying resolutions. For this evaluation, we utilized E9.5 mouse embryo data from the Stereo-seq platform (4,356 spots) and the Slide-seq V2 platform (14,758 spots) (Fig. 4B). The results revealed that stMSA effectively integrated the unbalanced slices, producing accurate joint domain identification results (Fig. 4B, C). In contrast, SEDR exhibited significant batch effects in its latent representation, as shown in the batch UMAP visualization, and STAligner struggled to identify joint domains (Supplementary Fig. S8A).

To further validate the reliability of stMSA’s domain identification results, we compared the spatial distribution of identified domains with the expression patterns of corresponding marker genes, which have been confirmed by previous biological experiments. Notable tissue structures, including the brain, heart, spinal cord, lung, and sclerotome, demonstrated a strong correlation with their respective marker genes: *Crabp2* for the brain [46], *Actc1* for the heart [47], *Map1b* for the spinal cord [48], *Ptn* for the lung [49], and *Pax1* for the sclerotome [50] (Supplementary Fig. S8B). These findings further corroborate the accuracy of stMSA in domain identification, demonstrating its applicability across various sequencing platforms and dataset sizes.

### 3.5 Deciphering B cell development trajectories through spatial multi-omics integration in human tonsils

To explore the integration of spatial proteomics and transcriptomics data, we applied stMSA to a human tonsil dataset, utilizing spatial-CITE-seq for proteomics and the 10X Visium platform for transcriptomics [3]. This integration enables a more comprehensive understanding of cellular functions [51], particularly for B cell development in the tonsil. Notably, the datasets exhibited significant differences in Unique Molecular Identifier (UMI) count distributions across spots, posing challenges for effective data harmonization (Fig. 4E).

stMSA demonstrated its capability to produce an expressive latent representation, as evidenced by Geary’s C and Moran’s I metrics. The lowest Geary’s C score and highest Moran’s I score achieved by stMSA indicate its effectiveness in capturing spatial patterns (Fig. 4F). Furthermore, stMSA successfully identified joint domains between RNA and protein slices, highlighting its robust performance in domain detection (Fig. 4G). stMSA integrated the spots from the two omics datasets more effectively than SEDR and STAligner. While SEDR assigned the brown domain 5 in the RNA slice and the green domain 3 in the protein slice, and STAligner assigned the purple domain 2 and the red domain 1, stMSA correctly identified and assigned domain 0 (the blue domain) to this region (Supplementary Fig. S9). This underscores stMSA’s superior ability to harmonize data patterns from heterogeneous sources in the human tonsil dataset.

To enhance domain resolution, we adjusted the Louvain clustering algorithm’s resolution parameter to 1.0, enabling finer domain delineation. This adjustment allowed us to clearly identify the lymph follicular region, with the follicular center designated as domain 6 and the extra-follicular plasma cell region as domain 3, aligning well with histological observations (Fig. 4H, Supplementary Fig. S10A). Subsequent analysis of spatially variable genes (SVGs) within the follicular region revealed associations with lymph follicular functions [52, 53].

Utilizing the latent representation generated by stMSA for trajectory inference, we found that identified domains aligned well with the developmental stages of B cells. Specific marker proteins and genes, such as *IgD* and *S100A8* for naive B cells [54], *IgD* and *IGHD* for mature B cells, and *IgM* and *CXCL12* for germinal center (GC) B cells [55, 56], corroborated this alignment (Fig. 4I). The spatial distribution of these markers closely corresponded with the domains identified by stMSA, showcasing a coherent relationship with B cell development. We also utilized CellChat [57] to analyze cell-cell communication based on domain identification results. The analysis revealed significant interactions among domains 2, 3, and 4, while domain 0 primarily interacted with domain 1. Notably, domain 1 engaged with all other domains, supporting the inferred B cell development trajectory. Pathway analysis indicated that these interactions were pathway-specific: domain 0 interacted with domain 1 via the EGF pathway, while domains 2, 3, and 4 primarily connected through the PDGF and CXCL pathways (Fig. 4J).

In summary, the experimental results demonstrate that stMSA is capable of integrating multi-omics data effectively. It successfully captures the developmental patterns within the multi-omics datasets, accurately identifies domain results, and constructs a correct developmental trajectory of B cells in the human tonsil dataset.

### 3.6 Elucidating tissue identity through cross-slice matching in embryonic development

Matching spots across adjacent spatial omics slices is critical for understanding the developmental trajectories of various cell types, tissues, and the process of organogenesis. This matching involves establishing correspondences between spot pairs from different slices taken at varying developmental stages, ensuring that matched spots exhibit similar gene expression profiles while preserving the spatial relationships inherent in each SO slice. To evaluate the cross-slice matching capabilities of stMSA, we applied the model to the Mouse Organogenesis Spatiotemporal Transcriptomic Atlas (MOSTA) dataset [58].

The results of applying stMSA to match tissue types between two mouse embryo slices collected on embryonic days 14.5 (E14.5) and 15.5 (E15.5) are illustrated in Fig. 5A. The Sankey plot in Fig. 5B demonstrates effective matching between these slices, revealing that most tissue types in mature organs, including the brain, heart, muscle, liver, and lung, have been accurately aligned. Furthermore, this matching process enabled us to identify specific organogenesis stages, such as the finding that cartilage at stage E15.5 develops from cartilage and cartilage primordium at stage E14.5. This developmental relationship is supported by the matching results shown in Fig. 5C, which highlights only the matched spot pairs between the two tissues. To further validate this relationship, we visualized the spatial distribution of *Col2a1*, a marker gene for cartilage [59], alongside the domain distribution of cartilage and cartilage primordium at stage E14.5. Notably, the expression pattern of *Col2a1* aligned closely with the corresponding domain distributions.

**Fig. 5.**
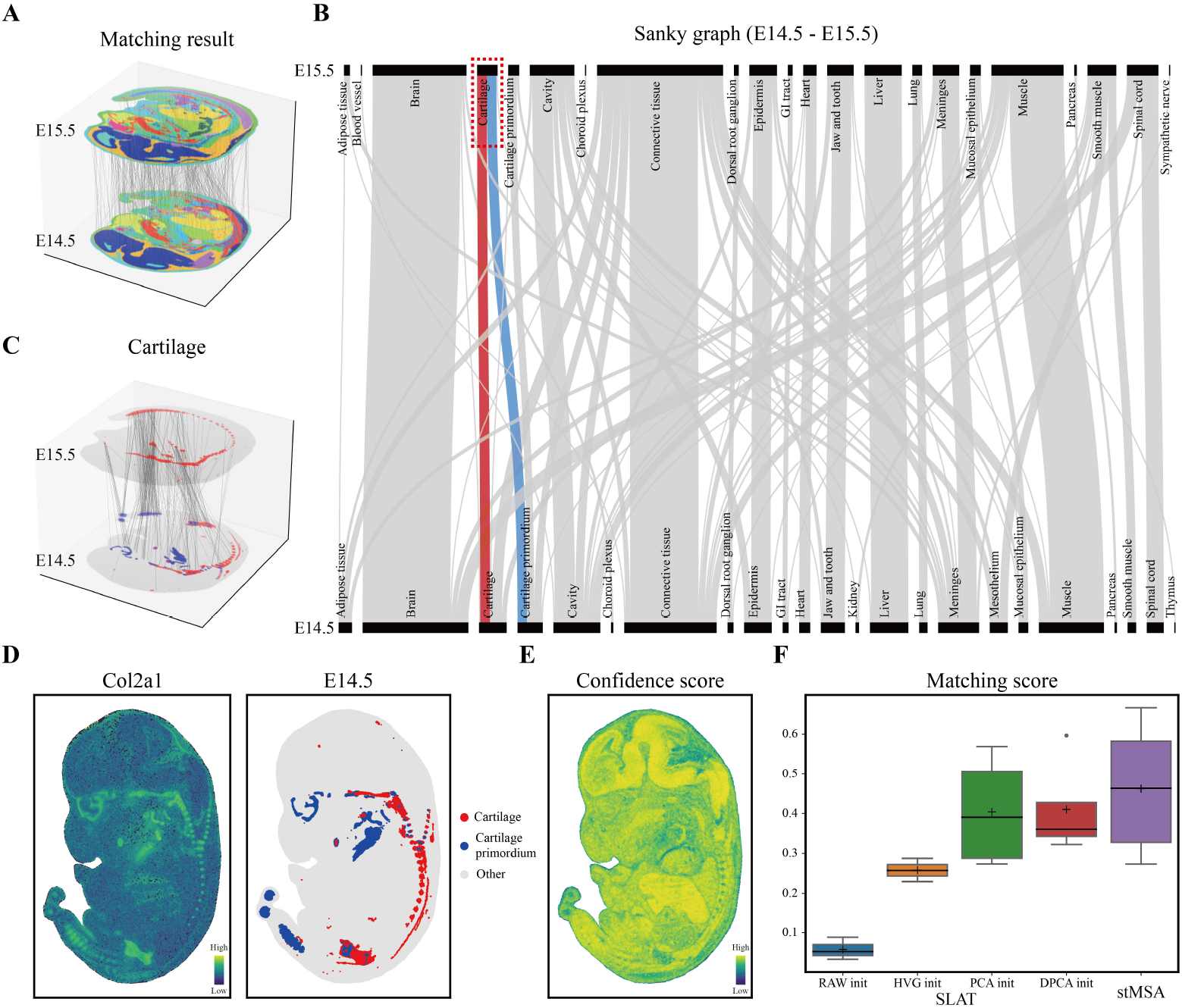
stMSA demonstrates effective tissue matching abilities in the Mouse Organogenesis Spatiotemporal Transcriptomic Atlas (MOSTA) dataset. **(A)** Visualization of the cross-slice matching results between E14.5 and E15.5 stages of mouse embryo. To improve the clarity of the matching results, 500 matched spot pairs were randomly selected. **(B)** The Sankey graph illustrating the tissue matching between E14.5 and E15.5 stages, with a highlighted path showcasing the matching relation between cartilage and cartilage primordium. **(C)** Matched spot pairs specifically focusing on the cartilage-cartilage primordium relationship at the E14.5 and E15.5 stages. **(D)** Marker gene expression patterns and spatial domain distributions of cartilage and cartilage primordium at E14.5 stage. **(E)** Confidence scores of the matching results, where higher scores indicate greater matching confidence, and vice versa. **(F)** Comparison of the matching scores between stMSA and SLAT under different settings (RAW, HVG, PCA, and DPCA init).

To quantitatively assess the overall matching performance of stMSA, we calculated a confidence score that evaluates the similarity between the embedded spots that were matched. This score, defined as the cosine distance between the embeddings of two matched spots, is presented in Fig. 5E for the E14.5 slice, clearly illustrating a high degree of similarity between the matched spots. Additionally, we extended our analysis to examine cross-slice matching between pairs at other stages, specifically E9.5-E10.5, E10.5-E11.5, E11.5-E12.5, E12.5-E13.5, and E13.5-E14.5. We compared the matching results with those obtained using the Spatial-linked Analysis of Transcriptomics (SLAT), which has four different initialization settings based on input types: using all genes (Raw init), only highly variable genes (HVG init), dimension-reduced gene expression profiles based on principal component analysis (PCA init), and dual PCA (DPCA init) [34]. Fig. 5F summarizes the matching scores obtained by stMSA and the four SLAT versions (detailed in Supplementary Note S1), demonstrating that stMSA outperforms SLAT across various settings.

### 3.7 stMSA enables the multi-slice alignment in mouse brain coronal sections

Current sequencing technologies limit researchers to capturing two-dimensional (2D) slices of biological tissues, posing challenges in fully understanding three-dimensional (3D) biological structures. To tackle this, we applied the stMSA method for automatic spatial omics data alignment on a dataset comprising seven coronal sections from mouse brains [60]. Notably, this dataset contains two flipped orientations labeled “Orient 1” and “Orient 2,” which can influence the integration and alignment of the slices.

For the alignment process, stMSA identifies a landmark domain to guide the Iterative Closest Point (ICP) algorithm in learning a rotation matrix (refer to Materials and Methods). As shown in Fig. 6B, stMSA initially identified spatial domains that aligned well with the Allen Brain Atlas annotations (Fig. 3E). Taking the pyramidal layer (domain 7) in the hippocampus as an example, this domain aligned seamlessly with the hook-like structure observed in the histology image (Fig. 6A) and the expression pattern of the marker gene *Hpca* [61] (Fig. 6C). This observation led to the selection of the pyramidal layer as the landmark domain for guiding the rotation matrix learning.

**Fig. 6.**
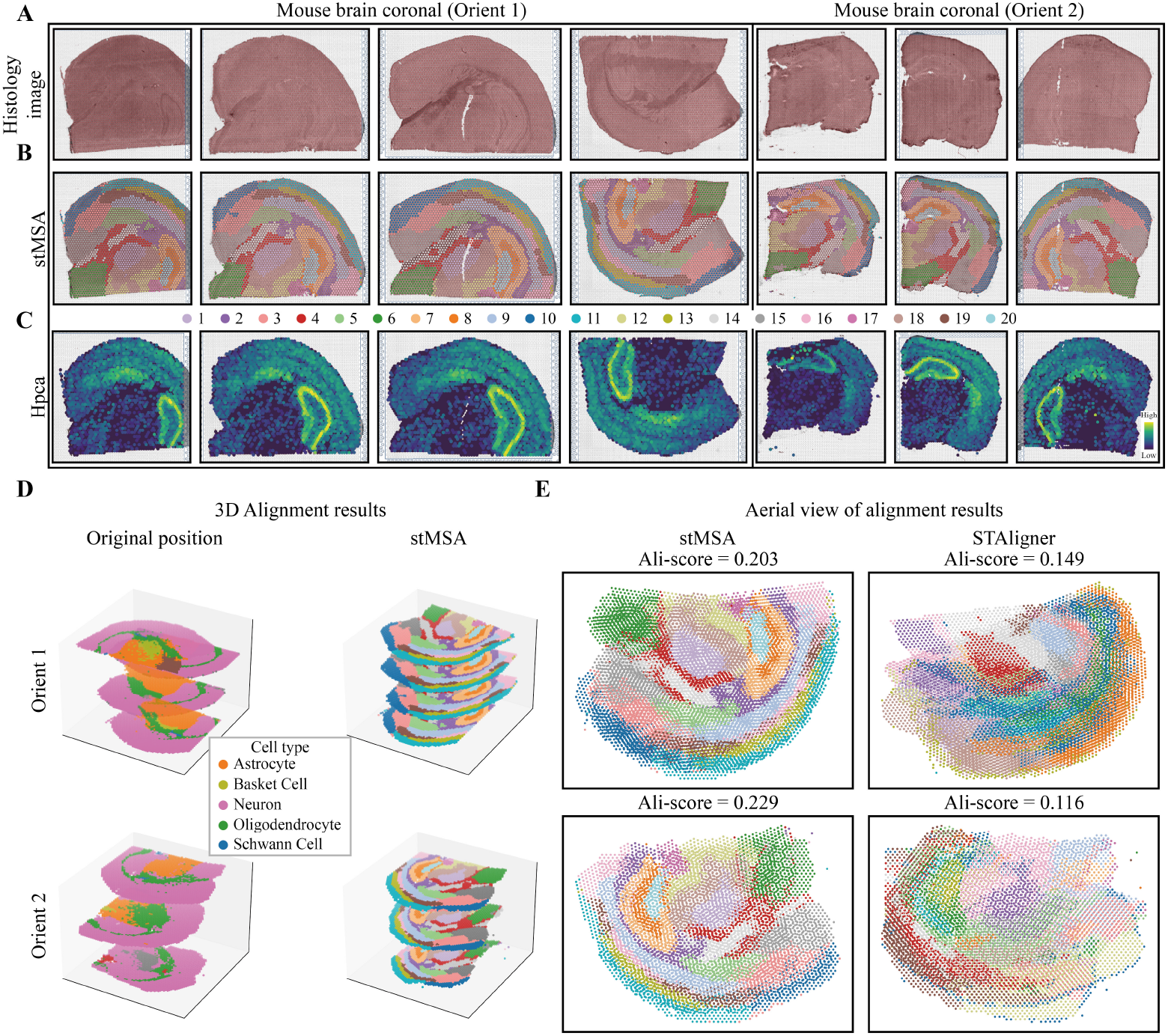
The performance of stMSA on a mouse brain coronal section SO dataset. **(A)** Original orientation of the histology image of the mouse brain coronal dataset. **(B)** Visualization of spatial domains detected by stMSA. **(C)** Spatial distribution of marker genes for the pyramidal layer in the hippocampus. **(D)** 3D visualization showing the original position of the dataset and the alignment results achieved by stMSA. **(E)** Aerial view of alignment results comparing stMSA with STAligner.

The challenges encountered during the alignment process are illustrated in Fig. 6D (left), which showcases the original positions and orientations of the dataset, revealing noticeable misalignment. To address this, we employed a modified ICP algorithm (refer to Materials and Methods) on the “Orient 1” and “Orient 2” datasets. This approach enabled stMSA to effectively coordinate the rotation and flipping of individual slices, seamlessly integrating them into a cohesive positional system. The alignment process ensures a uniform orientation across all slices, facilitating well-aligned visualization and interpretation of the spatial transcriptomics data in 3D views (Fig. 6D, right).

To evaluate stMSA’s alignment performance further, we compared it with STAligner. The aerial views of the aligned slices are presented in Fig. 6E. stMSA successfully aligns the slices, resulting in notable coherence in the proximity of spots belonging to the same domain. This visual coherence is depicted in the aerial views, which show domain distributions consistent with those observed in individual slices. In contrast, the distribution produced by STAligner appears less coherent. Additionally, we calculated an alignment score (see Materials and Methods) to quantitatively assess the alignment performance, demonstrating that stMSA outperforms STAligner in both orientations.

## 4 Discussion

The rapid advancement of sequencing technologies has resulted in the generation of an increasing volume of high-quality spatial omics data. This growth has created a pressing need to integrate spatial omics data from diverse sources, which may vary in experimental protocols, sequencing platforms, size, shape, and orientation. In this work, we present a novel contrastive learning-based method, termed stMSA, designed for the integration of multiple spatial omics datasets. stMSA learns cross-batch spot embeddings by optimizing two key principles: (1) enhancing the similarity of embeddings within a slice for spots situated in connected microenvironments and (2) maximizing the similarity of embeddings between cross-batch spots with analogous gene expression profiles. By adhering to these principles, stMSA effectively captures the distribution patterns of spots both within and across slices. Additionally, stMSA employs graph attention-based autoencoder techniques and deep embedding clustering to optimize model parameters and reconstruct gene expression profiles.

We thoroughly evaluate the performance of stMSA across a variety of downstream tasks, including batch-effect correction, joint domain detection, cross-slice matching, and multi-slice alignment. Our results demonstrate that stMSA offers a comprehensive solution for integrating multiple spatial omics slices from different sources, facilitating advanced analyses of spatial omics data. We strongly believe that as spatial omics sequencing technologies continue to evolve, stMSA will become an invaluable tool for researchers, empowering them to uncover novel biological insights from an increasingly diverse array of datasets.

## 5 Availability of data and materials

The implemented code is available online at https://github.com/hannshu/stMSA.

## 7 Acknowledgments

We thank all of the contributors of the open-source datasets and freely available tools used in this study. We also appreciate the suggestive comments of reviewers. This work has been supported by the National Natural Science Foundation of China (grant number 62102319) and the Natural Science Project of Shaanxi Provincial Department of Education (grant number 23JK0562).

## References

[1] Williams, C. G., Lee, H. J., Asatsuma, T., Vento-Tormo, R. & Haque, A. An introduction to spatial transcriptomics for biomedical research. Genome Medicine 14, 68 (2022). URL 10.1186/s13073-022-01075-1.

[2] Asp, M., Bergenstråhle, J. & Lundeberg, J. Spatially Resolved Transcriptomes—Next Generation Tools for Tissue Exploration. BioEssays 42, 1900221 (2020). URL https://onlinelibrary.wiley.com/doi/abs/10.1002/bies.201900221. eprint: https://onlinelibrary.wiley.com/doi/pdf/10.1002/bies.201900221.

[3] Liu, Y. et al. High-plex protein and whole transcriptome co-mapping at cellular resolution with spatial CITE-seq. Nature Biotechnology 41, 1405–1409 (2023). URL https://www.nature.com/articles/s41587-023-01676-0. Publisher: Nature Publishing Group.

[4] Cheng, M. et al. Spatially resolved transcriptomics: a comprehensive review of their technological advances, applications, and challenges. Journal of Genetics and Genomics 50, 625–640 (2023). URL https://www.sciencedirect.com/science/article/pii/S1673852723000759.

[5] Dong, K. & Zhang, S. Deciphering spatial domains from spatially resolved transcriptomics with an adaptive graph attention auto-encoder. Nature Communications 13, 1739 (2022). URL https://www.nature.com/articles/s41467-022-29439-6.

[6] Li, J., Chen, S., Pan, X., Yuan, Y. & Shen, H.-B. Cell clustering for spatial transcriptomics data with graph neural networks. Nature Computational Science 2, 399–408 (2022). URL https://www.nature.com/articles/s43588-022-00266-5.

[7] Schott, M. et al. Open-ST: High-resolution spatial transcriptomics in 3D (2023). URL https://www.biorxiv.org/content/10.1101/2023.12.22.572554v1. xPages: 2023.12.22.572554 Section: New Results.

[8] Hie, B., Bryson, B. & Berger, B. Efficient integration of heterogeneous single-cell transcriptomes using Scanorama. Nature Biotechnology 37, 685–691 (2019). URL https://www.nature.com/articles/s41587-019-0113-3.

[9] Korsunsky, I. et al. Fast, sensitive and accurate integration of single-cell data with Harmony. Nature Methods 16, 1289–1296 (2019). URL https://www.nature.com/articles/s41592-019-0619-0. Number: 12 Publisher: Nature Publishing Group.

[10] Yue, L. et al. A guidebook of spatial transcriptomic technologies, data resources and analysis approaches. Computational and Structural Biotechnology Journal 21, 940–955 (2023). URL https://www.sciencedirect.com/science/article/pii/S2001037023000156.

[11] Xu, H. et al. Unsupervised spatially embedded deep representation of spatial transcriptomics. Genome Medicine 16, 12 (2024). URL 10.1186/s13073-024-01283-x.

[12] Zhou, X., Dong, K. & Zhang, S. Integrating spatial transcriptomics data across different conditions, technologies and developmental stages. Nature Computational Science (2023). URL https://www.nature.com/articles/s43588-023-00528-w.

[13] Wang, G. et al. Construction of a 3D whole organism spatial atlas by joint modelling of multiple slices with deep neural networks. Nature Machine Intelligence 1–14 (2023). URL https://www.nature.com/articles/s42256-023-00734-1. Publisher: Nature Publishing Group.

[14] Wolf, F. A., Angerer, P. & Theis, F. J. SCANPY: large-scale single-cell gene expression data analysis. Genome Biology 19, 15 (2018). URL 10.1186/s13059-017-1382-0.

[15] Maynard, K. R. et al. Transcriptome-scale spatial gene expression in the human dorsolateral prefrontal cortex. Nature Neuroscience 24, 425–436 (2021). URL https://www.nature.com/articles/s41593-020-00787-0. Number: 3 Publisher: Nature Publishing Group.

[16] Jolliffe, I. T. & Cadima, J. Principal component analysis: a review and recent developments. Philosophical Transactions of the Royal Society A: Mathematical, Physical and Engineering Sciences 374, 20150202 (2016). URL https://royalsocietypublishing.org/doi/10.1098/rsta.2015.0202. Publisher: Royal Society.

[17] Veličković, P. et al. Graph Attention Networks (2018). URL http://arxiv.org/abs/1710.10903. 1710.10903 [cs, stat].

[18] Yosinski, J., Clune, J., Nguyen, A., Fuchs, T. & Lipson, H. Understanding Neural Networks Through Deep Visualization (2015). URL http://arxiv.org/abs/1506.06579. 1506.06579 [cs].

[19] Clevert, D.-A., Unterthiner, T. & Hochreiter, S. Fast and Accurate Deep Network Learning by Exponential Linear Units (ELUs) (2016). URL http://arxiv.org/abs/1511.07289. 1511.07289 [cs].

[20] Xie, J., Girshick, R. & Farhadi, A. Unsupervised Deep Embedding for Clustering Analysis (2016). URL http://arxiv.org/abs/1511.06335. 1511.06335 [cs].

[21] Krizhevsky, A., Sutskever, I. & Hinton, G. E. ImageNet classification with deep convolutional neural networks. Communications of the ACM 60, 84–90 (2017). URL https://dl.acm.org/doi/10.1145/3065386.

[22] Blondel, V. D., Guillaume, J.-L., Lambiotte, R. & Lefebvre, E. Fast unfolding of communities in large networks. Journal of Statistical Mechanics: Theory and Experiment 2008, P10008 (2008). URL http://arxiv.org/abs/0803.0476. 0803.0476 [cond-mat, physics:physics].

[23] Scrucca, L., Fop, M., Murphy, T. B. & Raftery, A. E. mclust 5: Clustering, Classification and Density Estimation Using Gaussian Finite Mixture Models. The R Journal 8, 289–317 (2016). URL https://journal.r-project.org/archive/2016/RJ-2016-021/index.html.

[24] Yeung, K. Y. & Ruzzo, W. L. Details of the Adjusted Rand indexand Clustering algorithms Supplement to the paper “An empirical study on Principal Component Analysis for clustering gene expression data” (to appear in Bioinformatics).

[25] Pedregosa, F. et al. Scikit-learn: Machine Learning in Python. Journal of Machine Learning Research 12, 2825–2830 (2011). URL http://jmlr.org/papers/v12/pedregosa11a.html.

[26] Jeffers, J. N. R. A Basic Subroutine for Geary’s Contiguity Ratio. Journal of the Royal Statistical Society. Series D (The Statistician) 22, 299–302 (1973). URL https://www.jstor.org/stable/2986827. Publisher: [Royal Statistical Society, Wiley].

[27] Moran, P. A. P. Notes on Continuous Stochastic Phenomena. Biometrika 37, 17–23 (1950). URL https://www.jstor.org/stable/2332142. Publisher: [Oxford University Press, Biometrika Trust].

[28] Douze, M. et al. The Faiss library (2024). URL https://arxiv.org/abs/2401.08281v1.

[29] Besl, P. & McKay, N. D. A method for registration of 3-D shapes. IEEE Transactions on Pattern Analysis and Machine Intelligence 14, 239–256 (1992). URL http://ieeexplore.ieee.org/document/121791/.

[30] Yang, J., Li, H., Campbell, D. & Jia, Y. Go-ICP: A Globally Optimal Solution to 3D ICP Point-Set Registration. IEEE Transactions on Pattern Analysis and Machine Intelligence 38, 2241–2254 (2016). URL http://arxiv.org/abs/1605.03344. 1605.03344 [cs].

[31] Kingma, D. P. & Ba, J. Adam: A Method for Stochastic Optimization (2017). URL http://arxiv.org/abs/1412.6980. 1412.6980 [cs].

[32] Fey, M. & Lenssen, J. E. Fast Graph Representation Learning with PyTorch Geometric (2019). URL http://arxiv.org/abs/1903.02428. 1903.02428 [cs, stat].

[33] Xu, H. et al. SPACEL: deep learning-based characterization of spatial transcriptome architectures. Nature Communications 14, 7603 (2023). URL https://www.nature.com/articles/s41467-023-43220-3. Number: 1 Publisher: Nature Publishing Group.

[34] Xia, C.-R., Cao, Z.-J.Tu, X.-M. & Gao, G. Spatial-linked alignment tool (SLAT) for aligning heterogenous slices. Nature Communications 14, 7236 (2023). URL https://www.nature.com/articles/s41467-023-43105-5. Number: 1 Publisher: Nature Publishing Group.

[35] Xu, Z. et al. STOmicsDB: a comprehensive database for spatial transcriptomics data sharing, analysis and visualization. Nucleic Acids Research 52, D1053– D1061 (2024). URL 10.1093/nar/gkad933.

[36] Wei, S. et al. Charting the Spatial Transcriptome of the Human Cerebral Cortex at Single-Cell Resolution (2024). URL https://www.biorxiv.org/content/10.1101/2024.01.31.576150v1. Pages: 2024.01.31.576150 Section: New Results.

[37] Wolf, F. A. et al. PAGA: graph abstraction reconciles clustering with trajectory inference through a topology preserving map of single cells. Genome Biology 20, 59 (2019). URL 10.1186/s13059-019-1663-x.

[38] Sullivan, K. E. et al. Sharp cell-type-identity changes differentiate the retrosplenial cortex from the neocortex. Cell Reports 42, 112206 (2023). URL https://www.sciencedirect.com/science/article/pii/S2211124723002176.

[39] Grimstvedt, J. S. et al. A multifaceted architectural framework of the mouse claustrum complex. Journal of Comparative Neurology 531, 1772– 1795 (2023). URL https://onlinelibrary.wiley.com/doi/abs/10.1002/cne.25539. eprint: https://onlinelibrary.wiley.com/doi/pdf/10.1002/cne.25539.

[40] Hoerder-Suabedissen, A. et al. Temporal origin of mouse claustrum and development of its cortical projections. Cerebral Cortex (New York, NY) 33, 3944–3959 (2022). URL https://www.ncbi.nlm.nih.gov/pmc/articles/PMC10068282/.

[41] Fulcher, B. D., Murray, J. D., Zerbi, V. & Wang, X.-J. Multimodal gradients across mouse cortex. Proceedings of the National Academy of Sciences of the United States of America 116, 4689–4695 (2019). URL https://www.ncbi.nlm.nih.gov/pmc/articles/PMC6410879/.

[42] Sandebring-Matton, A. et al. Microdissected Pyramidal Cell Proteomics of Alzheimer Brain Reveals Alterations in Creatine Kinase B-Type, 14-3-3-gamma, and Heat Shock Cognate 71. Frontiers in Aging Neuroscience 13 (2021). URL https://www.frontiersin.org/articles/10.3389/fnagi.2021.735334.

[43] Wei, X. et al. Single-cell Stereo-seq reveals induced progenitor cells involved in axolotl brain regeneration. Science 377, eabp9444 (2022). URL https://www.science.org/doi/10.1126/science.abp9444. Publisher: American Association for the Advancement of Science.

[44] Stickels, R. R. et al. Highly sensitive spatial transcriptomics at near-cellular resolution with Slide-seqV2. Nature Biotechnology 39, 313–319 (2021). URL https://www.nature.com/articles/s41587-020-0739-1. Number: 3 Publisher: Nature Publishing Group.

[45] Ståhl, P. L. et al. Visualization and analysis of gene expression in tissue sections by spatial transcriptomics. Science 353, 78–82 (2016). URL https://www.science.org/doi/10.1126/science.aaf2403. Publisher: American Association for the Advancement of Science.

[46] Pietilä, R. et al. Molecular anatomy of adult mouse leptomeninges. Neuron 111, 3745–3764.e7 (2023). URL https://www.cell.com/neuron/abstract/S0896-6273(23)00666-9. Publisher: Elsevier.

[47] Frank, D. et al. Cardiac α-Actin (ACTC1) Gene Mutation Causes Atrial-Septal Defects Associated With Late-Onset Dilated Cardiomyopathy. Circulation: Genomic and Precision Medicine (2019). URL https://www.ahajournals.org/doi/10.1161/CIRCGEN.119.002491. Publisher: Lippincott Williams & Wilkin-sHagerstown, MD.

[48] Tortosa, E. et al. Microtubule-associated Protein 1B (MAP1B) Is Required for Dendritic Spine Development and Synaptic Maturation. Journal of Biological Chemistry 286, 40638–40648 (2011). URL https://www.sciencedirect.com/science/article/pii/S0021925820504158.

[49] Weng, T. & Liu, L. The role of pleiotrophin and β-catenin in fetal lung development. Respiratory Research 11, 80 (2010). URL 10.1186/1465-9921-11-80.

[50] Dong, J. et al. Single-cell RNA-seq analysis unveils a prevalent epithelial/mesenchymal hybrid state during mouse organogenesis. Genome Biology 19, 31 (2018). URL 10.1186/s13059-018-1416-2.

[51] Lundberg, E. & Borner, G. H. H. Spatial proteomics: a powerful discovery tool for cell biology. Nature Reviews Molecular Cell Biology 20, 285–302 (2019). URL https://www.nature.com/articles/s41580-018-0094-y. Publisher: Nature Publishing Group.

[52] Hwang, I.-Y., Hwang, K.-S., Park, C., Harrison, K. A. & Kehrl, J. H. Rgs13 Constrains Early B Cell Responses and Limits Germinal Center Sizes. PLoS ONE 8, e60139 (2013). URL https://www.ncbi.nlm.nih.gov/pmc/articles/PMC3606317/.

[53] Boyd, S. D., Natkunam, Y., Allen, J. R. & Warnke, R. A. Selective Immunophenotyping for Diagnosis of B-cell Neoplasms: Immunohistochemistry and Flow Cytometry Strategies and Results. Applied immunohistochemistry & molecular morphology : AIMM /official publication of the Society for Applied Immunohistochemistry 21, 116–131 (2013). URL https://www.ncbi.nlm.nih.gov/pmc/articles/PMC4993814/.

[54] Kitagori, K. et al. Expression of S100A8 protein on B cells is associated with disease activity in patients with systemic lupus erythematosus. Arthritis Research & Therapy 25, 76 (2023). URL 10.1186/s13075-023-03057-z.

[55] Barinov, A. et al. Essential role of immobilized chemokine CXCL12 in the regulation of the humoral immune response. Proceedings of the National Academy of Sciences of the United States of America 114, 2319–2324 (2017). URL https://www.ncbi.nlm.nih.gov/pmc/articles/PMC5338526/.

[56] Morgan, D. & Tergaonkar, V. Unraveling B cell trajectories at single cell resolution. Trends in Immunology 43, 210–229 (2022). URL https://www.cell.com/t rends/immunology/abstract/S1471-4906(22)00003-5. Publisher: Elsevier.

[57] Jin, S. et al. Inference and analysis of cell-cell communication using CellChat. Nature Communications 12, 1088 (2021). URL https://www.nature.com/articles/s41467-021-21246-9. Number: 1 Publisher: Nature Publishing Group.

[58] Chen, A. et al. Spatiotemporal transcriptomic atlas of mouse organogenesis using DNA nanoball-patterned arrays. Cell 185, 1777–1792.e21 (2022). URL https://www.cell.com/cell/abstract/S0092-8674(22)00399-3. Publisher: Elsevier.

[59] Hissnauer, T. N. et al. Identification of molecular markers for articular cartilage. Osteoarthritis and Cartilage 18, 1630–1638 (2010). URL https://www.sciencedirect.com/science/article/pii/S1063458410003328.

[60] Ballester Roig, M. N. et al. Probing pathways by which rhynchophylline modifies sleep using spatial transcriptomics. Biology Direct 18, 21 (2023). URL 10.1186/s13062-023-00377-7.

[61] Tzingounis, A. V., Kobayashi, M., Takamatsu, K. & Nicoll, R. A. Hippocalcin Gates the Calcium Activation of the Slow Afterhyperpolarization in Hippocampal Pyramidal Cells. Neuron 53, 487–493 (2007). URL https://www.sciencedirect.com/science/article/pii/S0896627307000311.

