## Supplementary for "Efficient integration of spatial omics data for joint domain detection, matching, and alignment with stMSA"

#### Supplementary Notes

##### Supplementary Note S1 Comparison with other state-of-the-art methods

For multi-slice integration and domain identification, we compared stMSA against several leading methods: Scanorama[1], Harmony[2], STAGATE[3], SEDR[4], STAligner[5], Stitch3D[6], and SPACEL[7]. For cross-slice matching, our comparison involved stMSA and SLAT[8]. Additionally, for multi-slice alignment, we directly compared stMSA with STAligner. Below, we provide introductions and list parameter settings for these methods:

**Scanorama** effectively recognizes datasets that contain cells with similar transcriptional profiles and utilizes these matches for batch correction and data integration. Importantly, it achieves this without merging non-overlapping datasets.

We utilized Scanorama in its default configuration and employed the *scanorama.correct_scanpy()* function for data integration. Additionally, we applied K-Means[9] clustering to identify multi-slice domains using embeddings generated by Scanorama. The source code of Scanorama can be found at https://github.com/brianhie/scanorama.

**Harmony** consists of two primary steps: the clustering step and the batch correction step. In the clustering step, embeddings are used to generate soft cluster results. These results are then utilized in the batch correction step to produce new, batch-corrected embeddings. Through iterative processing, Harmony ultimately outputs the batch-corrected embeddings of the input datasets.

We employed the Python implementation of Harmony, known as Harmonypy, using the *harmonypy.run_harmony()* function for data integration. Harmony was operated with its default settings. For clustering, we applied the K-Means method to the embeddings generated by Harmony. The source code for Harmony, implemented in Python, is available at https://github.com/slowkow/harmonypy.

**STAGATE** employs a graph attention autoencoder network to learn latent embeddings. This is achieved by maximizing the similarity between input gene expressions and their reconstructed outputs from the decoder.

We utilized the PyTorch Geometric implementation of STAGATE, running it with the default settings. For clustering, we employed the Mclust clustering method[10], which is also the default clustering method offered by STAGATE. The source code for STAGATE is available at https://github.com/QIFEIDKN/STAGATE_pyG.

**SEDR** generates latent representations for multi-slice datasets using a graph variational autoencoder. Subsequently, it corrects batch effects using the Harmony algorithm.

We applied SEDR as instructed in the documentation located at https://sedr.readthedocs.io/en/latest/Tutorial3_Batch_integration.html. For input, we used 200 principal components (PCs) obtained through the PCA algorithm[11], which were then processed by SEDR. Following this, we ran Harmony to obtain batch-corrected embeddings from those learned by SEDR. For clustering, we used the Louvain[12] method for the mouse olfactory dataset and the Mclust method for other datasets. The source code for SEDR is available at https://github.com/JinmiaoChenLab/SEDR.

**STAligner** utilizes a graph attention autoencoder to generate latent representations. The model parameters are optimized by maximizing the similarity between input gene expressions and the reconstructed gene expressions outputted by the autoencoder. Additionally, the model optimizes the similarity of positive Mutually Nearest Neighbor (MNN) pairs while minimizing the similarity of negative MNN pairs.

We employed STAligner as per the instructions provided at https://staligner.readthedocs.io/en/latest/. We used the script available in the instructions and conducted clustering as recommended, utilizing the Louvain method for the mouse olfactory dataset and the Mclust method for other datasets. The source code for STAligner is available at https://github.com/zhoux85/STAligner.

**Stitch3D** processes single-cell reference data and multi-slice ST data as inputs. It constructs a 3D spatial graph for the ST data and trains latent representations using a graph attention network. Additionally, Stitch3D optimizes the model parameters by refining the estimation of cell type proportions and mitigating slice-spot and slice-gene specific effects.

We utilized Stitch3D following the instructions provided at https://stitch3d-tutorial.readthedocs.io/en/latest/tutorials/DLPFC/STitch3D_DLPFC.html. We used the single-cell reference data provided by Stitch3D and generated clustering results using the Gaussian Mixture Model[13] as recommended. The source code for Stitch3D can be found at https://github.com/YangLabHKUST/STitch3D.

**SPACEL** first uses single-cell data as a reference to generate deconvolution results for spatial transcriptomics data. It then uses the resulting proportion matrix as input to create latent embeddings for each spot using a graph convolutional network.

We utilized SPACEL, applying it with the single-cell data it provided. It is worth noticed that the single-cell data SPACEL used is differ from the one Stitch3D used. We followed the testing script at https://spacel.readthedocs.io/en/latest/tutorials/Visium_human_DLPFC_Spoint.html as instruction. The clustering results were obtained using the *splane_model.identify_spatial_domain()* function. The source code for SPACEL is available at https://github.com/QuKunLab/SPACEL.

**SLAT** employs a lightweight graph convolutional autoencoder network to learn embeddings between two slices. The model parameters are optimized by maximizing the Wasserstein distance[14] between the embeddings of the two slices and enhancing the similarity between the input gene expressions and their reconstructed outputs from the autoencoder. The SLAT has four initialization settings based on the type of inputs: using all genes (Raw init), using the highly variable genes (HVG init), using dimension-reduced gene expression profile based on principle component analysis (PCA init) or dual-principal component analysis (DPCA init).

We utilized SLAT by following the script provided in its documentation at https://slat.readthedocs.io/en/latest/tutorials/basic_usage.html. The source code for SLAT is available at https://github.com/gao-lab/SLAT.

### Supplementary Figures

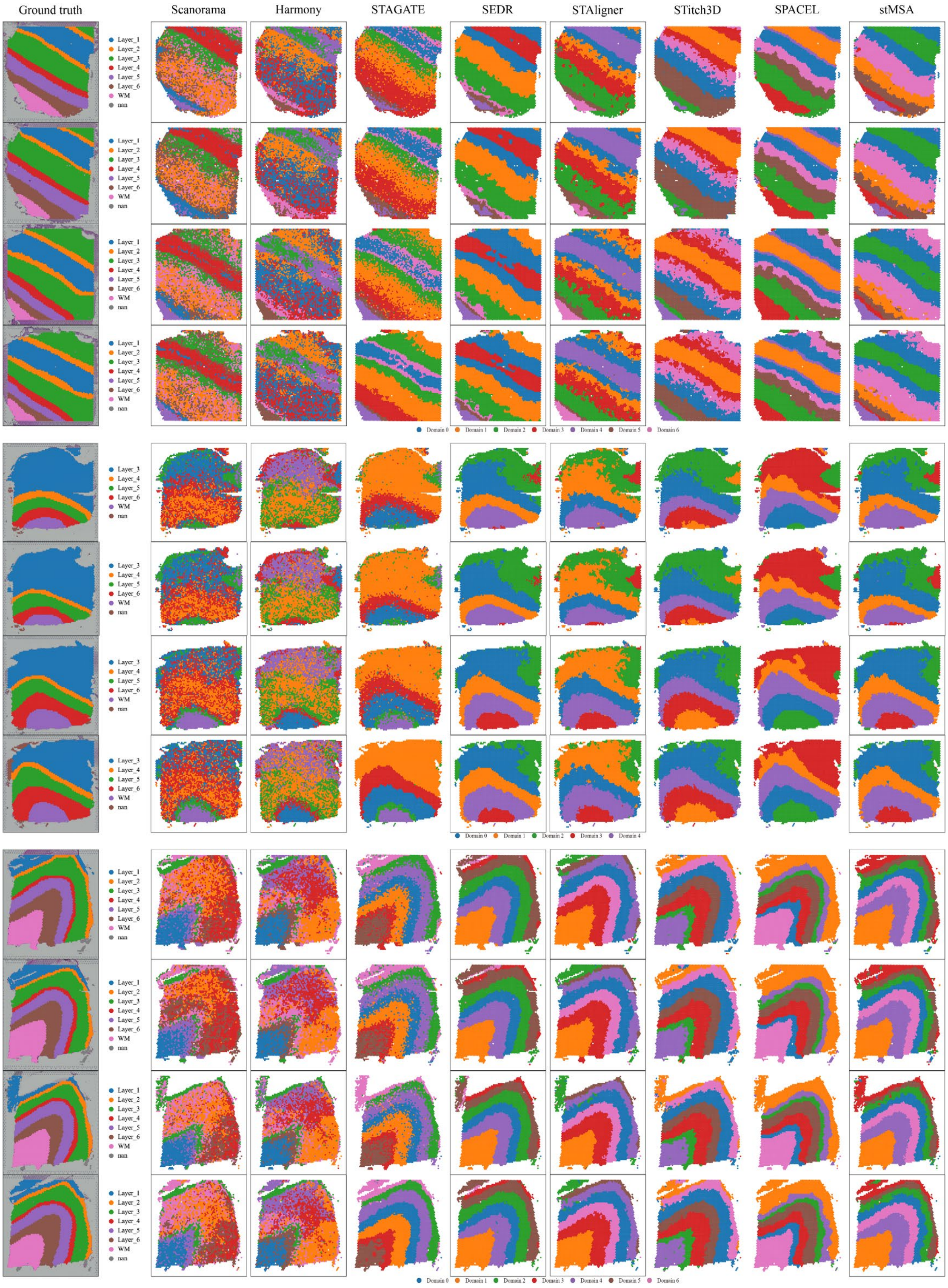

**Supplementary Figure S1. The spatial domain identification results of the four slices from three donors using different state-of-the-art methods.**

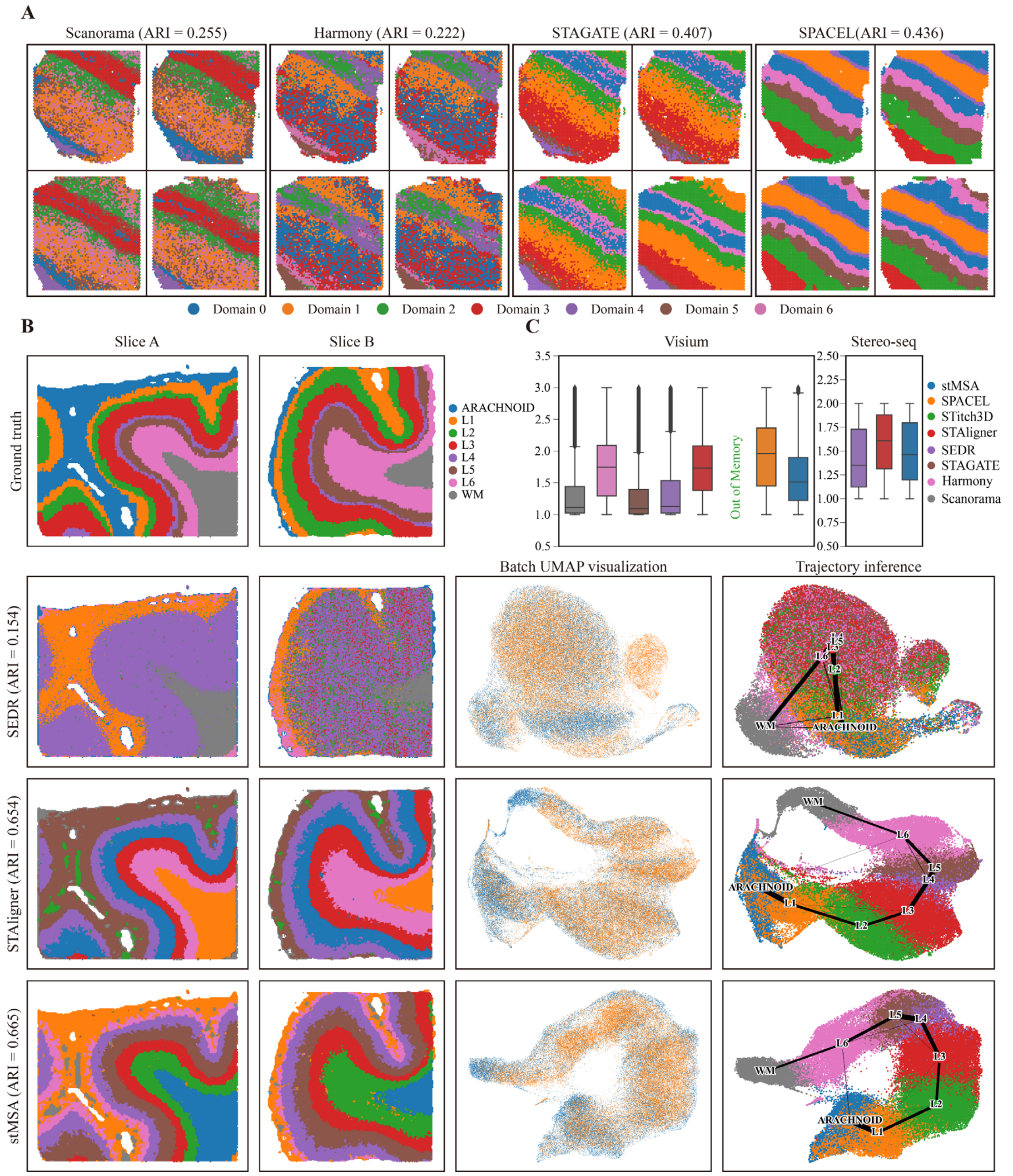

**Supplementary Figure S2. The domain identification result for the 10X Visium DLPFC dataset, and the domain identification result for Stereo-seq DLPFC data. (A)** The domain identification result for donor 1 for Scanorama, Harmony, STAGATE, and Stitch3D in the 10X Visium DLPFC dataset. **(B)** The ground truth annotation, the domain identification result, the batch UMAP visualization, and the trajectory inference result for Stereo-seq DLPFC dataset for SEDR, STAligner, and stMSA. **(C)** The LISI batch mixture score for 10X Visium and Stereo-seq obtained DLPFC dataset. In this context, the term *batch* refers to different donors for the 10X Visium DLPFC dataset and two distinct tissue slices for the Stereo-seq DLPFC dataset. Stitch3D is unable to process all 12 slices due to the excessive demand for GPU memory in the 10X Visium DLPFC dataset.

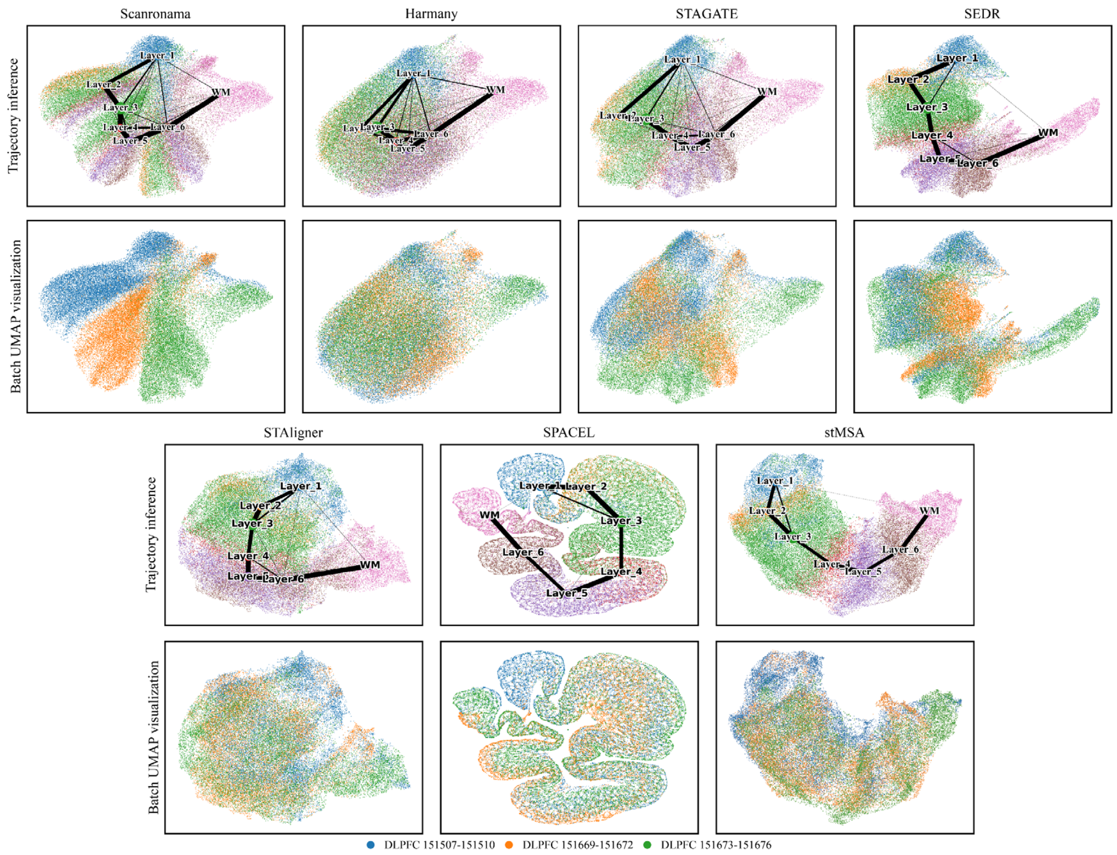

**Supplementary Figure S3. The UMAP plot of the embeddings generated by six methods for all 12 slices**. In each method, the upper plot showcases the UMAP plot colored by domain label and trajectory inference result lied on it, while the lower UMAP plot is colored by the donor label.

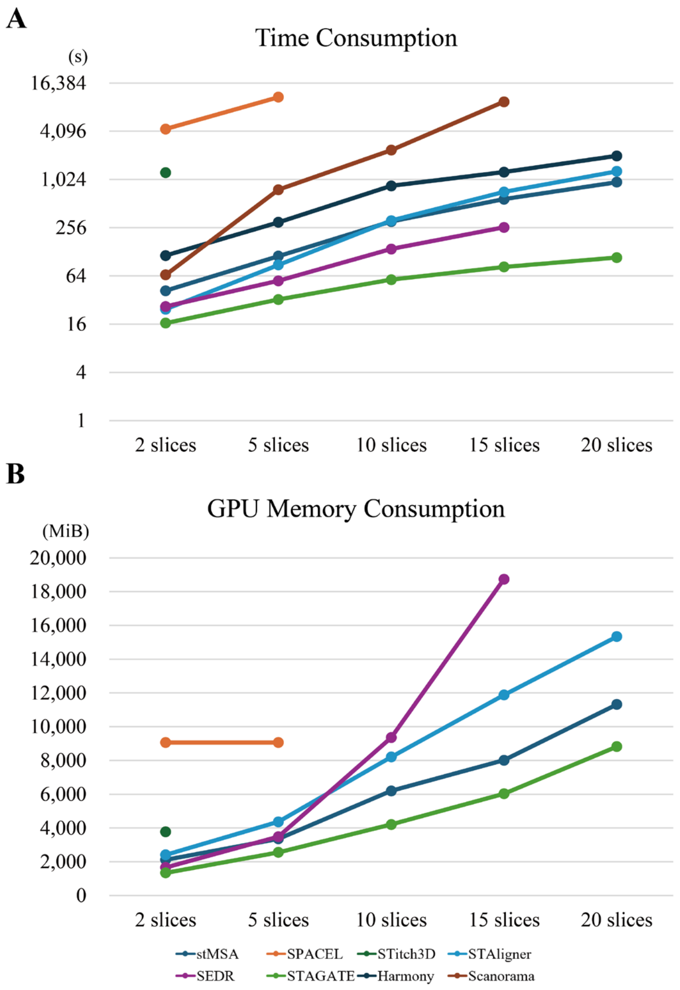

**Supplementary Figure S4. The summary of resource consumption of the seven methods in DLPFC 151507 slice. (A)** The time consumption of the seven methods for representation learning on 2, 5, 10, 15, and 20 DLPFC 151507 slice. **(B)** The GPU memory consumption of the five methods for representation learning on 2, 5, 10, 15, and 20 DLPFC 151507 slice. Harmony and Scanorama do not use GPU resource for representation learning.

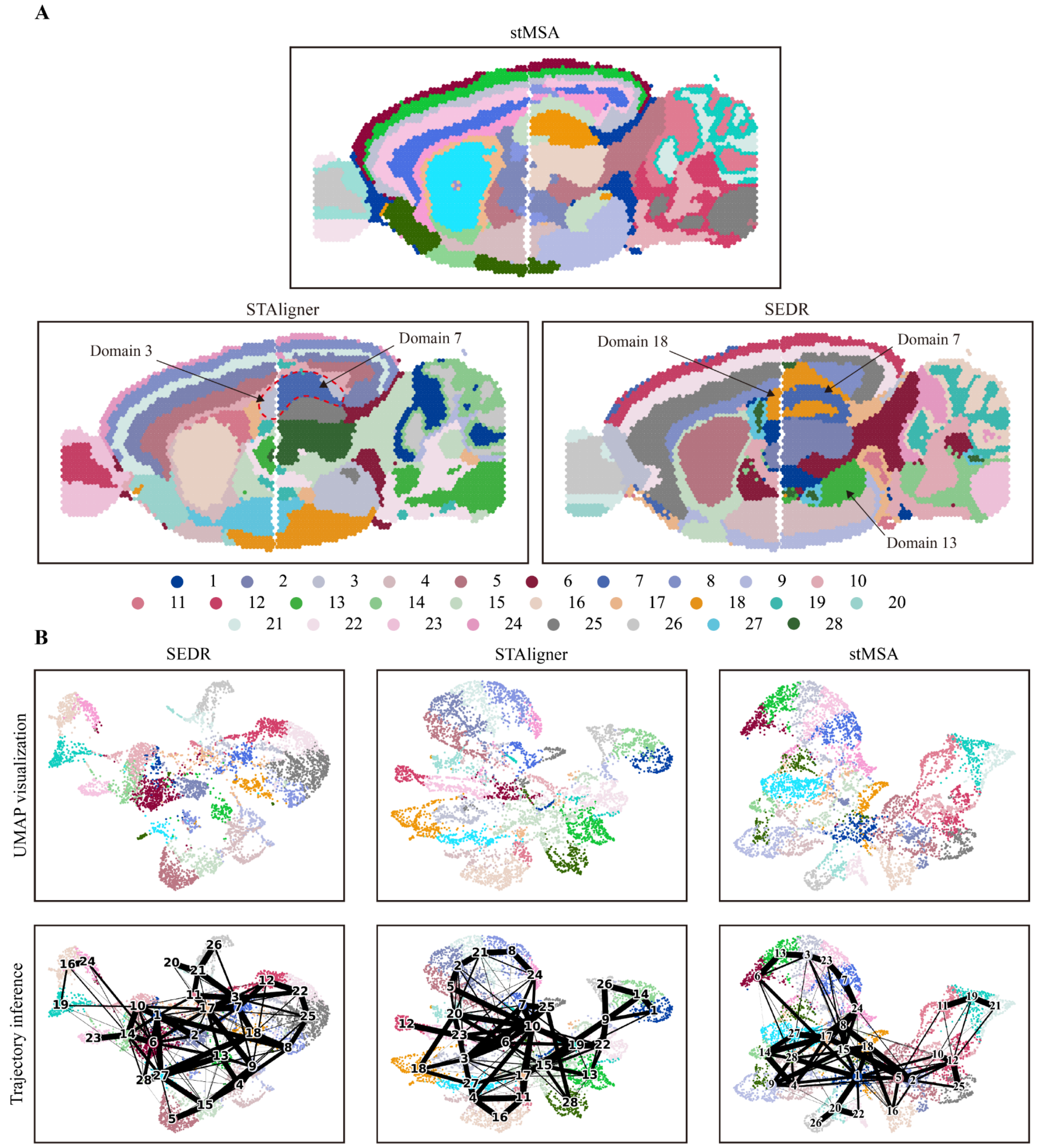

**Supplementary Figure S5. The domain identification and UMAP plot of a mouse brain sagittal section. (A)** The domain identification result achieved by stMSA, STAligner, and SEDR. **(B)** The UMAP plot and the trajectory inference plot of the learned latent representation by three methods, colored by predicted label. The UMAP plot of stMSA demonstrates a similar relative position to their actual position in the isocortex region.

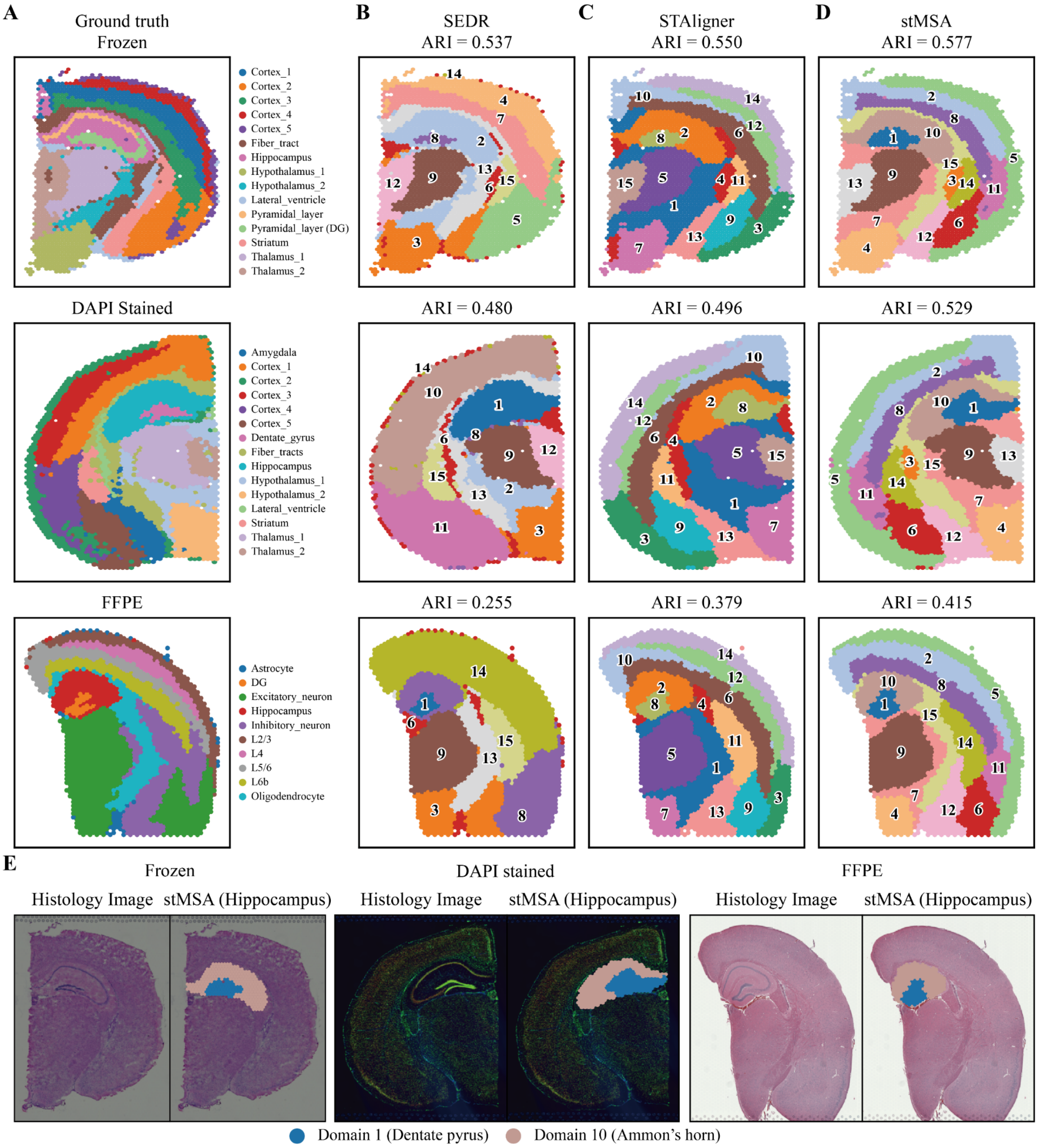

**Supplementary Figure S6. The domain identification results of coronal mouse brain section dataset.** The manual annotations **(A)** and domain detection results by SEDR **(B)**, STAligner **(C)**, and stMSA **(D)** across mouse brain coronal sections derived from frozen slices, DAPI-stained slices, and FFPE slices, respectively. **(E)** The histology image and the hippocampus region predicted by stMSA in frozen, DAPI stained, and FFPE mouse brain coronal section, respectively.

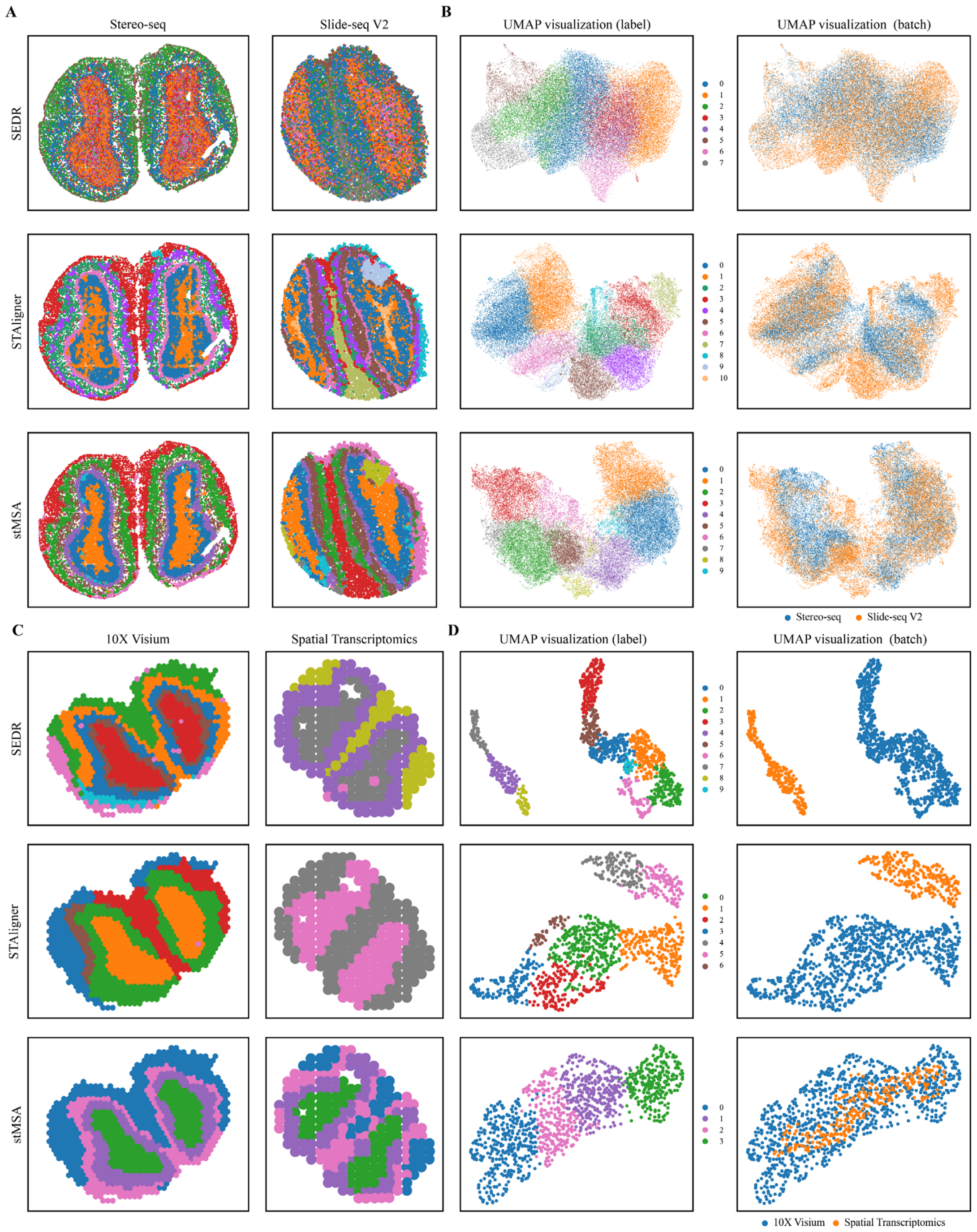

**Supplementary Figure S7. The domain identification result and UMAP visualization of the Stereo-seq Slide-seq V2, and 10X Visium Spatial Transcriptomcis obtained mouse olfactory bulb for SEDR, STAligner, and stMSA, respectively. (A)** The domain identification results of the Stereo-seq Slide-seq V2 dataset of SEDR, STAligner, and stMSA, respectively. **(B)** The UMAP visualization results of the Stereo-seq Slide-seq V2 dataset of SEDR, STAligner, and stMSA, respectively. **(C)** The domain identification results of the 10X Visium Spatial Transcriptomics dataset of SEDR, STAligner, and stMSA, respectively. **(D)** The UMAP visualization results of the 10X Visium Spatial Transcriptomics dataset of SEDR, STAligner, and stMSA, respectively.

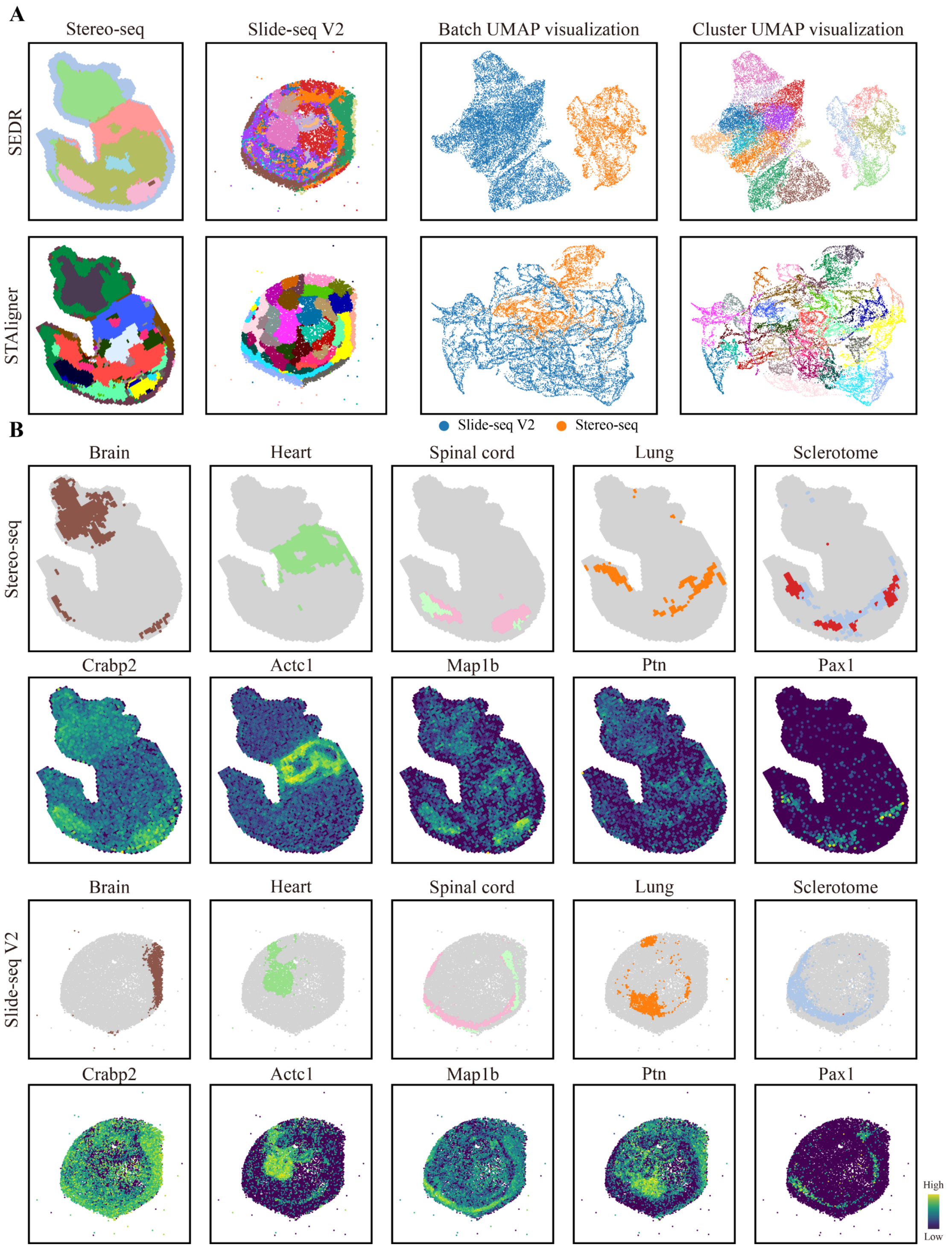

**Supplementary Figure S8. The domain identification result and spatial distribution of the tissue structure and the corresponding marker gene for the mouse embryo dataset. (A)** The domain identification results and UMAP visualization results of the mouse embryo dataset of SEDR and STAligner. **(B)** The brain, heart, spinal cord, lung, and sclerotome domain structure identified by stMSA and their corresponding marker gene for Stereo-seq and Slide-seq V2 obtained mouse embryo.

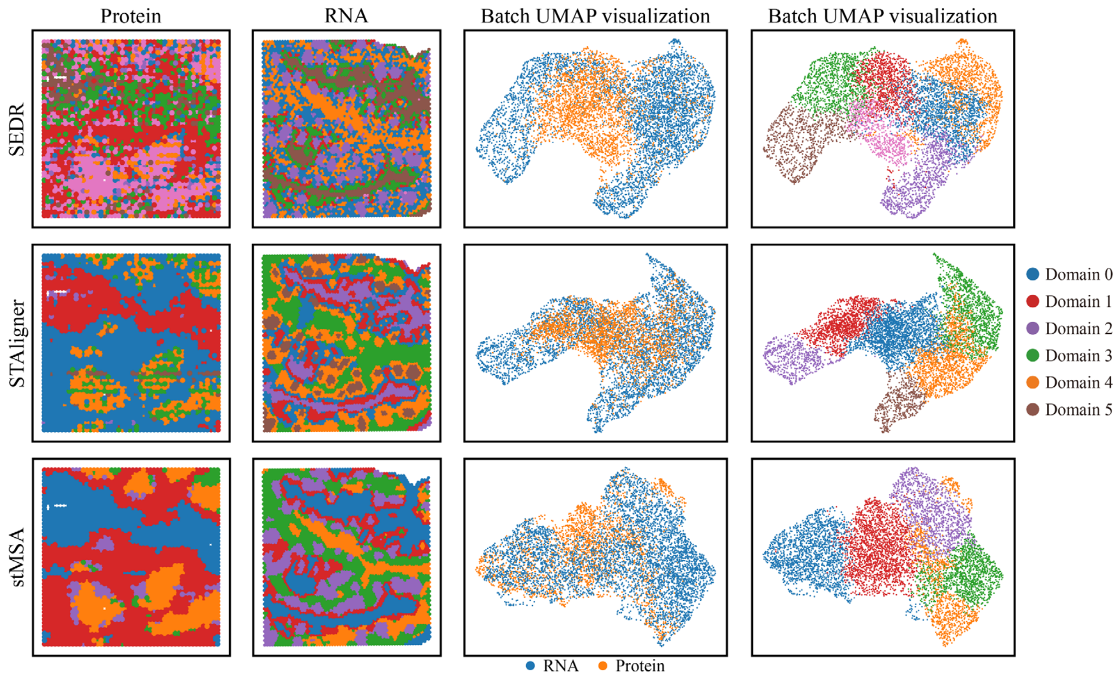

**Supplementary Figure S9. The domain identification result and batch UMAP visualization for the RNA-Protein human tonsil dataset.**

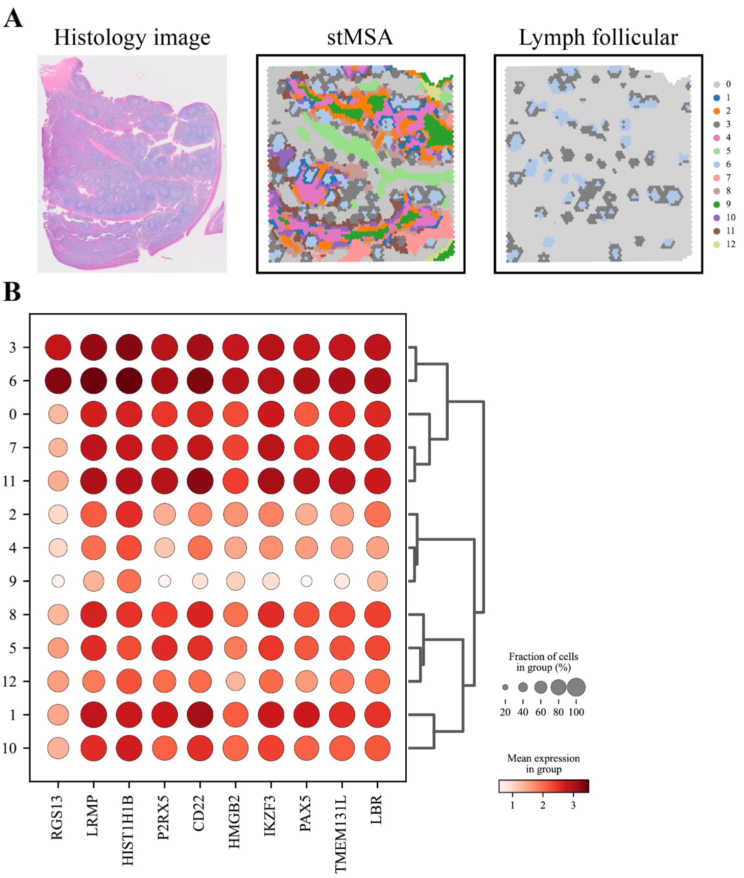

**Supplementary Figure S10. The high-resolution domain identification result and the spatial variable genes for the lymph follicular region. (A)** The histology image and the high-resolution domain identification results of the transcriptomics tonsil slice. **(B)** The spatial variable genes for the lymph follicular region.

#### Supplementary Tables

**Supplementary Table S1. Summary of all spatial transcriptomic datasets used for experiments in this study.**

<table>
<tr><th>Tissue</th><th>Section ID</th><th>Number of spots</th><th>Number of genes</th><th>Number of clusters</th><th>Sequencing technology</th><th>Label source</th></tr>
<tr><td rowspan="14">Human Brain Cortex</td><td>DLPFC 151507</td><td>4,226</td><td rowspan="12">33,538</td><td rowspan="4">7</td><td rowspan="12">10x Visium</td><td rowspan="12">Lieber Institute GitHub Repo</td></tr>
<tr><td>DLPFC 151508</td><td>4,384</td></tr>
<tr><td>DLPFC 151509</td><td>4,789</td></tr>
<tr><td>DLPFC 151510</td><td>4,634</td></tr>
<tr><td>DLPFC 151669</td><td>3,661</td><td rowspan="4">5</td></tr>
<tr><td>DLPFC 151670</td><td>3,498</td></tr>
<tr><td>DLPFC 151671</td><td>4,110</td></tr>
<tr><td>DLPFC 151672</td><td>4,015</td></tr>
<tr><td>DLPFC 151673</td><td>3,639</td><td rowspan="4">7</td></tr>
<tr><td>DLPFC 151674</td><td>3,673</td></tr>
<tr><td>DLPFC 151675</td><td>3,592</td></tr>
<tr><td>DLPFC 151676</td><td>3,460</td></tr>
<tr><td>Slice A</td><td>31,329</td><td>31,749</td><td rowspan="2">8</td><td rowspan="2">Stereo-seq</td><td rowspan="2">stMSA</td></tr>
<tr><td>Slice B</td><td>35,772</td><td>31,972</td></tr>
<tr><td rowspan="2">Mouse brain Saggital</td><td>Anterior</td><td>2,696</td><td rowspan="2">31,053</td><td rowspan="2">-</td><td rowspan="11">10x Visium</td><td rowspan="2">None</td></tr>
<tr><td>Posterior</td><td>3,353</td></tr>
<tr><td rowspan="9">Mouse brain Coronal</td><td>Frozen</td><td>2,688</td><td>18,078</td><td>15</td><td rowspan="2">squidpy built-in</td></tr>
<tr><td>DAPI Stained</td><td>2,800</td><td>16,562</td><td>15</td></tr>
<tr><td>FFPE</td><td>2,264</td><td>19,465</td><td>10</td><td>STOmicsDB</td></tr>
<tr><td>GSM6704280</td><td>2,522</td><td rowspan="6">32,285</td><td rowspan="6">-</td><td rowspan="6">None</td></tr>
<tr><td>GSM6704281</td><td>2,831</td></tr>
<tr><td>GSM6704282</td><td>2,752</td></tr>
<tr><td>GSM6704283</td><td>2,816</td></tr>
<tr><td>GSM6704284</td><td>2,108</td></tr>
<tr><td>GSM6704285</td><td>2,639</td></tr>
</table>

| | | | | | | |
|---|---|---|---|---|---|---|
| | GSM6704286 | 2,741 | | | | |
| Mouse olfactory bulb | - | 21,724 | 21,220 | | Slide-seq V2 | |
| | - | 19,109 | 27,106 | | Stereo-seq | |
| | - | 267 | 16,573 | | Spatial Transcriptomics | |
| | - | 918 | 31,053 | | 10x Visium | |
| Human Tonsil | - | 4,908 | 18,085 | | | |
| Mouse Embryo | - | 14,758 | 27,554 | | Slide-seq V2 | |
| | E9.5 E2S2 | 4,356 | 24,107 | 13 | Stereo-seq | STOmicsDB |
| | E9.5 E1S1 | 5,913 | 25,568 | 12 | | |
| | E10.5 E1S1 | 18,408 | 25,201 | 13 | | |
| | E11.5 E1S1 | 30,124 | 26,854 | 19 | | |
| | E12.5 E1S1 | 51,365 | 27,810 | 23 | | |
| | E13.5 E1S1 | 77,369 | 28,408 | 19 | | |
| | E14.5 E1S1 | 102,519 | 28,463 | 26 | | |
| | E15.5 E1S1 | 113,350 | 28,798 | 26 | | |

**Supplementary Table S2. Summary of spatial proteomics datasets used for experiments in this study**.

| Tissue | Number of spots | Number of proteins | Sequencing technology |
|---|---|---|---|
| Human Tonsil | 2,492 | 410 | spatial-CITE-seq |

#### Reference

[1] B. Hie, B. Bryson, and B. Berger, "Efficient integration of heterogeneous single-cell transcriptomes using Scanorama," *Nat. Biotechnol.*, vol. 37, no. 6, pp. 685–691, Jun. 2019, doi: 10.1038/s41587-019-0113-3.

[2] I. Korsunsky *et al.*, "Fast, sensitive and accurate integration of single-cell data with Harmony," *Nat. Methods*, vol. 16, no. 12, Art. no. 12, Dec. 2019, doi: 10.1038/s41592-019-0619-0.

[3] K. Dong and S. Zhang, "Deciphering spatial domains from spatially resolved transcriptomics with an adaptive graph attention auto-encoder," *Nat. Commun.*, vol. 13, no. 1, p. 1739, Apr. 2022, doi: 10.1038/s41467-022-29439-6.

[4] J. Chen *et al.*, "Unsupervised Spatially Embedded Deep Representation of Spatial Transcriptomics," In Review, preprint, Jul. 2021. doi: 10.21203/rs.3.rs-665505/v1.

[5] X. Zhou, K. Dong, and S. Zhang, "Integrating spatial transcriptomics data across different conditions, technologies and developmental stages," *Nat. Comput. Sci.*, Oct. 2023, doi: 10.1038/s43588-023-00528-w.

[6] G. Wang, J. Zhao, Y. Yan, Y. Wang, A. R. Wu, and C. Yang, "Construction of a 3D whole organism spatial atlas by joint modelling of multiple slices with deep neural networks," *Nat. Mach. Intell.*, pp. 1–14, Oct. 2023, doi: 10.1038/s42256-023-00734-1.

[7] H. Xu *et al.*, "SPACEL: deep learning-based characterization of spatial transcriptome architectures," *Nat. Commun.*, vol. 14, no. 1, Art. no. 1, Nov. 2023, doi: 10.1038/s41467-023-43220-3.

[8] C.-R. Xia, Z.-J. Cao, X.-M. Tu, and G. Gao, "Spatial-linked alignment tool (SLAT) for aligning heterogenous slices," *Nat. Commun.*, vol. 14, no. 1, Art. no. 1, Nov. 2023, doi: 10.1038/s41467-023-43105-5.

[9] J. MacQueen, "Some methods for classification and analysis of multivariate observations," *Proc. Fifth Berkeley Symp. Math. Stat. Probab. Vol. 1 Stat.*, vol. 5.1, pp. 281–298, Jan. 1967.

[10] L. Scrucca, M. Fop, T. B. Murphy, and A. E. Raftery, "mclust 5: Clustering, Classification and Density Estimation Using Gaussian Finite Mixture Models," *R J.*, vol. 8, no. 1, pp. 289–317, 2016.

[11] I. T. Jolliffe and J. Cadima, "Principal component analysis: a review and recent developments," *Philos. Trans. R. Soc. Math. Phys. Eng. Sci.*, vol. 374, no. 2065, p. 20150202, Apr. 2016, doi: 10.1098/rsta.2015.0202.

[12] V. D. Blondel, J.-L. Guillaume, R. Lambiotte, and E. Lefebvre, "Fast unfolding of communities in large networks," *J. Stat. Mech. Theory Exp.*, vol. 2008, no. 10, p. P10008, Oct. 2008, doi: 10.1088/1742-5468/2008/10/P10008.

[13] N. Kambhatla and T. Leen, "Classifying with Gaussian Mixtures and Clusters," in *Advances in Neural Information Processing Systems*, MIT Press, 1994. Accessed: May 30, 2023. [Online]. Available: https://proceedings.neurips.cc/paper_files/paper/1994/hash/621461af90cadfdaf0e8d4cc25129f91-Abstract.html

[14] M. Arjovsky, S. Chintala, and L. Bottou, "Wasserstein GAN." arXiv, Dec. 06, 2017. doi: 10.48550/arXiv.1701.07875.
